# PAannotate: Software for Exploring *Pseudomonas aeruginosa* Gene-Expression Associations from Transcriptomic Data

**DOI:** 10.64898/2026.11.24.768722

**Authors:** Robert W. Targ, Paul L. Bollyky

**Affiliations:** 300 Pasteur Dr., Rm L121, Division of Infectious Diseases and Geographic Medicine, Department of Medicine, Stanford University, Stanford, CA, USA. 93405

## Abstract

With the proliferation of transcriptomic data for *Pseudomonas aeruginosa* there is a need for software programs able to efficiently analyze both individual transcriptomes as well as large, publicly available compendiums of arrays. PAannotate is an R package and native macOS application capable of analyzing the Whiteley Lab database of 5,821 transcriptomes from 470 BioProjects. In the application, researchers can use recorded metadata such as strain, medium or treatment to select experiments or genes associated with particular expression patterns. Alternatively, selecting one gene of interest, a user can find other, coordinately regulated genes that move together across many environments, mutants, and treatments. PAannotate also allows users to select groups of genes and to use Gene Ontology and KEGG annotations and enrichment analysis to see whether those genes are associated with particular biological functions more often than expected by chance, determine whether a particular perturbation produces a reproducible response across studies, or find public experiments resembling your own experiment. Researchers can build and customize heatmaps and association networks. In one project, we have examined the *P. aeruginosa* transcriptomic associations with the genes encoding the Pf prophage present in *P. aeruginosa* strain PAO1. We found a strong negative correlation between PA0724, a Pf4 gene, and ccmG, a bacterial gene involved in energy metabolism. We identified that much of this relationship was reflected by a difference in whether the bacteria were grown with acetate or glycerol as their carbon source. This finding illustrates how PAannotate helps researchers investigate *P. aeruginosa* gene correlations.

## Introduction

When two genes rise and fall together in one experiment, do they still do so when the bacteria grow under different conditions? The question matters for *Pseudomonas aeruginosa*, a bacterium that can persist in chronic wounds and in the lungs of people with cystic fibrosis [56–58, 69]. RNA sequencing (RNA-seq) estimates how much RNA from each gene is present in a sample. The resulting profile, or transcriptome, records which genes the bacteria were expressing when they were sampled. Two genes may track one another because both respond to the growth medium. To judge whether their relationship extends beyond that experiment, a researcher needs to know which samples were compared and what happened in other studies.

Comparisons across experiments have already changed how microbiologists study this organism. Cornforth and colleagues measured bacterial RNA in human infections and found that genes involved in metabolism and communication between cells were expressed differently in patients and in laboratory cultures [69]. Lewin and colleagues in the Whiteley laboratory used patient sputum, the mucus coughed up from the lungs, as a reference for judging models of cystic fibrosis infection. Two laboratory models reproduced different parts of the patient expression profile. The public data also helped identify limited zinc availability as one reason a model differed from sputum [44]. Duncan and colleagues found that expression measurements in cystic fibrosis models were largely reproducible across laboratories, although the way samples were prepared for sequencing introduced additional variation [3].

Other investigators have asked what genes repeatedly appear together in these data. Tan and colleagues developed ADAGE, a method that learns recurring patterns from expression measurements without first assigning genes to pathways, the sets of reactions that carry out a biological task [39, 40]. A later version picked up a phosphate-starvation response even in growth media containing moderate amounts of phosphate [38]. Rajput and colleagues used independent component analysis, a statistical method that separates patterns that can vary independently across experiments. They found 104 groups of genes in 364 transcriptomes, and linked 81 groups to known regulators, which are molecules that control gene expression [42]. Such patterns can suggest which genes share a regulator. Establishing that one gene’s expression influences another requires additional evidence [42, 66, 67]. The iModulonDB resource lets readers examine these groups as well as relationships between individual genes under selected conditions [7, 37, 43].

Several resources make the experiments themselves available. The Gene Expression Omnibus (GEO) holds study descriptions and processed measurements, and its records link to the underlying sequencing reads in the Sequence Read Archive [70]. Doing and colleagues assembled public RNA-seq data into reusable tables of gene counts and supplied the code for updating them [1, 41]. Using those data, Lee and colleagues compared genes shared by two laboratory strains, PAO1 and PA14, with genes found in only one. Expression of strain-specific genes also helped them check how the samples had been labeled [2]. The Whiteley laboratory separately distributes gene counts and descriptions of the source samples and provides tools for searching and extracting them [8]. We have created a software program called “PAannotate’ to provide an accessible and rigorous method of exploring the Whiteley dataset. In the version we prepared for PAannotate, the collection comprises 5,821 sample records in 470 project groups (Fig. 1A).

**Figure 1.**
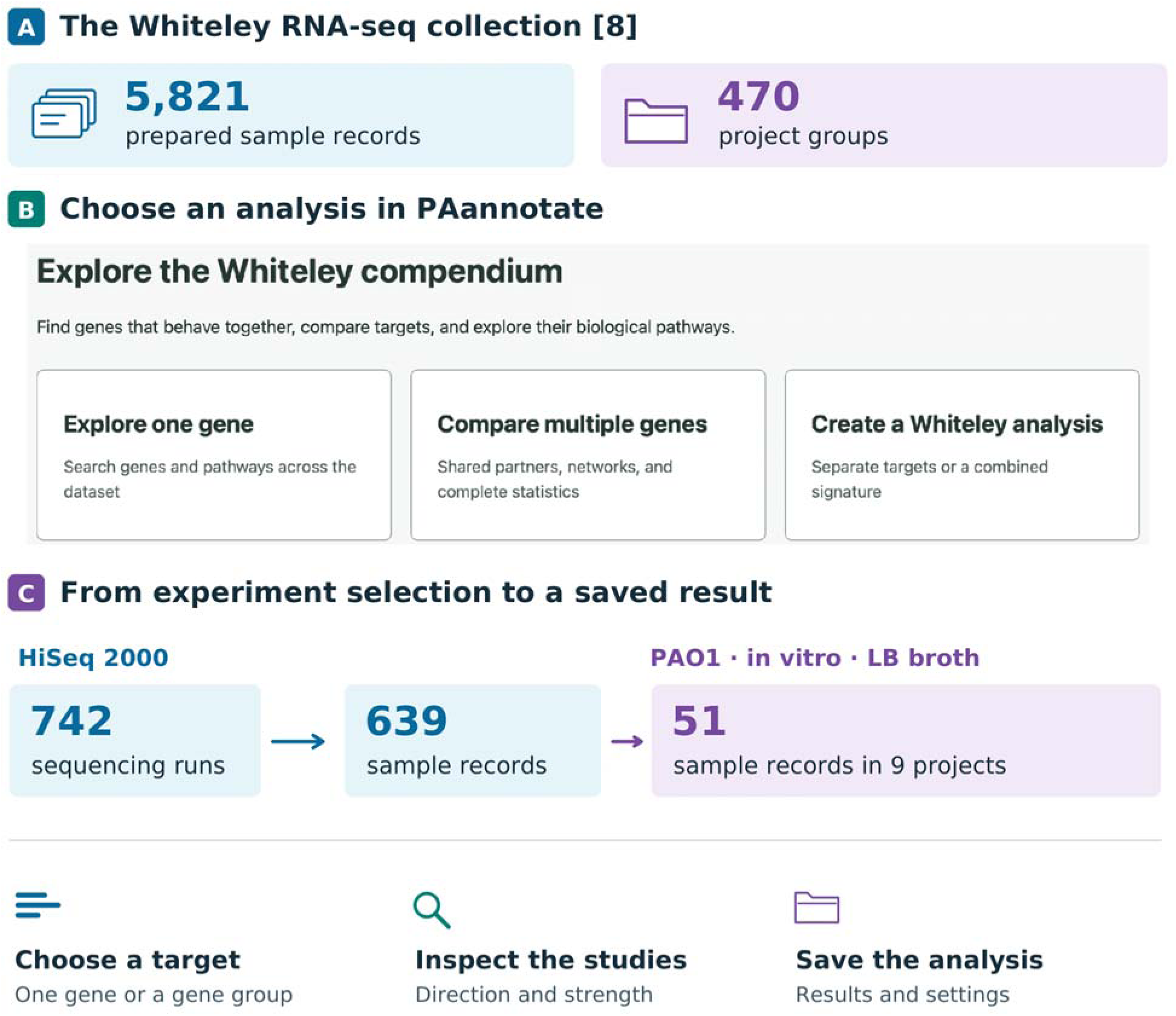
From a biological question to a saved PAannotate analysis. A) the Whiteley collection supplies RNA measurements and experimental descriptions [8]. B) a crop of the desktop home screen shows the three entry points. C) the recorded instrument search illustrates how several sequencing runs can represent one sample. Restricting the selection to PAO1 grown in vitro in LB broth yielded 51 records in nine projects. The lower row shows the choices available after cohort selection. Figure 4 illustrates a separate eight-sample search.

A microbiologist can already use COLOMBOS to compare a gene’s response across experimental conditions [4, 35], or GAPE and CF-Seq to find studies in which its expression changed between groups [5, 6]. The bacteria.guru database connects genes whose expression tracks together, and QIAGEN CLC Genomics Workbench imports gene counts and compares groups of samples [36, 71]. For a researcher starting from a particular gene in the Whiteley collection, a different question often follows: “Which other genes track it within the experiments relevant to my question, and how much does the answer depend on which experiments I include?” We developed PAannotate to connect that question of samples to the individual study results behind an association.

PAannotate combines a frontend macOS application with an analytical package written in R, a language used for statistics. Researchers can select experiments by recorded details such as strain or growth medium, choose one gene or a group, and examine which other genes vary in the same or the opposite direction. They can inspect the studies behind a result, look up what the associated genes do, and save the settings and results together (Fig. 1). We demonstrate that workflow in the application, use the R package and accompanying scripts for a broader comparison across studies, and test how accurately the software calculates associations and describes their uncertainty.

For the biological example, we chose Pf4, a thread-shaped (filamentous) virus that infects *P. aeruginosa*. A virus that infects bacteria is called a bacteriophage, or phage. Figure 2A shows Pf4 filaments and the protein coat that surrounds its DNA [30]. In a common laboratory strain of *P. aeruginosa*, PAO1, a copy of that Pf4 DNA occupies a part of the bacterial chromosome. This integrated stretch of viral DNA, called a prophage, spans 12,437 base pairs at positions 785,311– 797,747 in the PAO1 reference chromosome (NC_002516.2) [64, 65]. Figure 2C locates the eleven genes we could follow in the expression data, though the function of some of these genes, like PA0719, remains uncertain [29] (Fig. S7).

**Figure 2.**
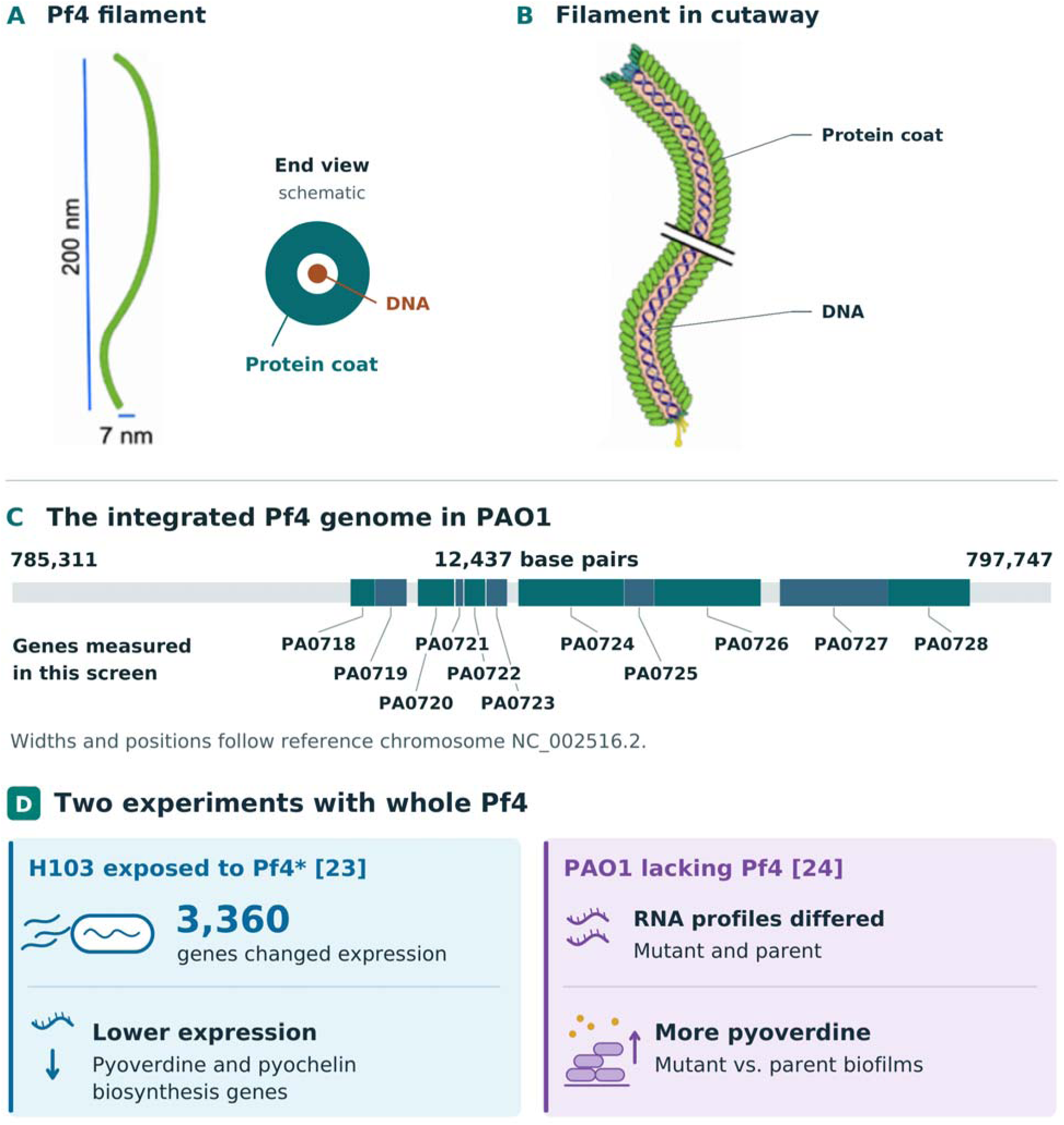
Pf4 particles, their place in the PAO1 chromosome, and their effects on the host. A) A schematic shows the threadlike Pf4 particle with the scale bars supplied in the drawing. The end view shows the protein coat surrounding the viral DNA. B) An enlarged cutaway shows the DNA inside the coat. The double slash marks a break in the drawing. C) colored segments locate the eleven measured genes within the integrated prophage [64, 65]. Gray denotes the remainder of the region. D) two published whole-phage experiments used different strains and interventions [23, 24]. Pyoverdine production in the deletion study was measured separately from RNA abundance.

In Pf, genes PA0718 and PA0728 work to insert viral genetic information into the bacterial genome. After integration, the bacterium performs its normal transcription of DNA into RNA, which now includes reproducing the viral genes as well. PA0720 forms a preliminary coat around a strand of newly-copied Pf DNA, and PA0721, PA0722, PA0724, and PA0725 act as caps that seal the DNA inside the protein coat. Before PA0726 helps get the new Pf4 virus out of the bacterial cell (without lysing it), PA0723 acts as a secondary, final coat protein that replaces what was created by PA0720. After all of that is done, the bacterium can release filamentous phage that may then go on to infect more bacteria [47]. Outside the cell, Pf particles have been shown to help organize biofilms, which are communities of bacteria embedded in material they produce, and they can form a protective sheath around individual cells [22, 30, 49, 60] (Fig. 2B). Pf delayed wound healing in animal experiments, and in people with cystic fibrosis, researchers found an association between Pf and chronic airway infection, and between high Pf levels in sputum and faster loss of lung function [56, 57, 58]. Laboratory work has also linked Pf to slower movement of airway mucus, reduced growth of cells that repair the airway lining, and slower movement of some antibiotics through sputum [50, 51, 59].

In *P. aeruginosa* strain H103, exposure to a Pf4 variant reduced production of pyocyanin- -a bacterial pigment--and activity of elastase, which is an enzyme that breaks down proteins [23]. Removing a resident Pf phage also affected traits linked to infection, with different effects in different strains [25].

Here, we used PAannotate to examine the *P. aeruginosa* transcriptomic associations with the genes encoding the Pf prophage. When their RNA levels rise or fall, do the same genes elsewhere in the bacterial chromosome tend to follow? Using the R package, we compared eleven locations, PA0718 through PA0728, in the PAO1 reference genome and compared the bacterial host genes associated with each of these 11 (Figs. 2C and 3). Related phages share some of these sequences, so RNA assigned to a reference location can have more than one possible source [45, 53]. We then traced one association back to the conditions in a single experiment and compared it with evidence from the wider collection (Figs. 4 and 5). Finally, we asked how the genes identified in the screen responded in two experiments set aside beforehand. Tortuel and colleagues exposed strain H103 to the Pf4* variant in a study deposited in GEO as GSE201738 [23, 62]. Meanwhile, Ismail and colleagues studied PAO1 after removal of its resident Pf4 [24]. Both experiments contribute counts to the Whiteley collection. They let us ask whether the selected genes, as a group, responded more strongly than other genes with similar baseline expression (Fig. 6).

**Figure 3.**
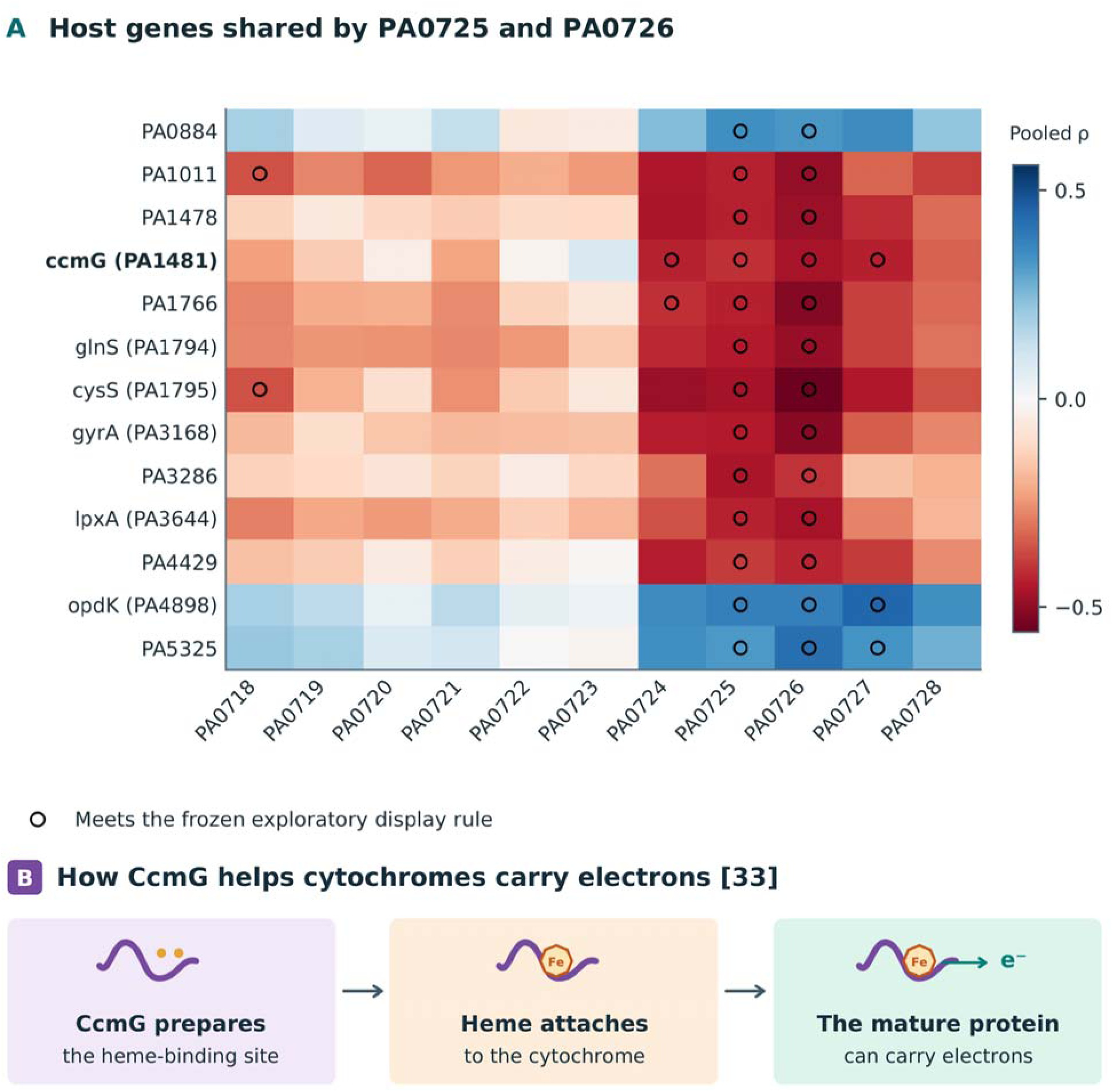
Host associations shared by PA0725 and PA0726. A) rows show the thirteen host genes that met the selection rule for both PA0725 and PA0726. Color gives the pooled Spearman correlation with each of the eleven Pf-region genes. Circles mark pairs meeting the exploratory selection rule in Methods. B) the known role of CcmG in preparing electron-carrying proteins [33] gives biological context to the highlighted row. The heatmap measures RNA associations.

**Figure 4.**
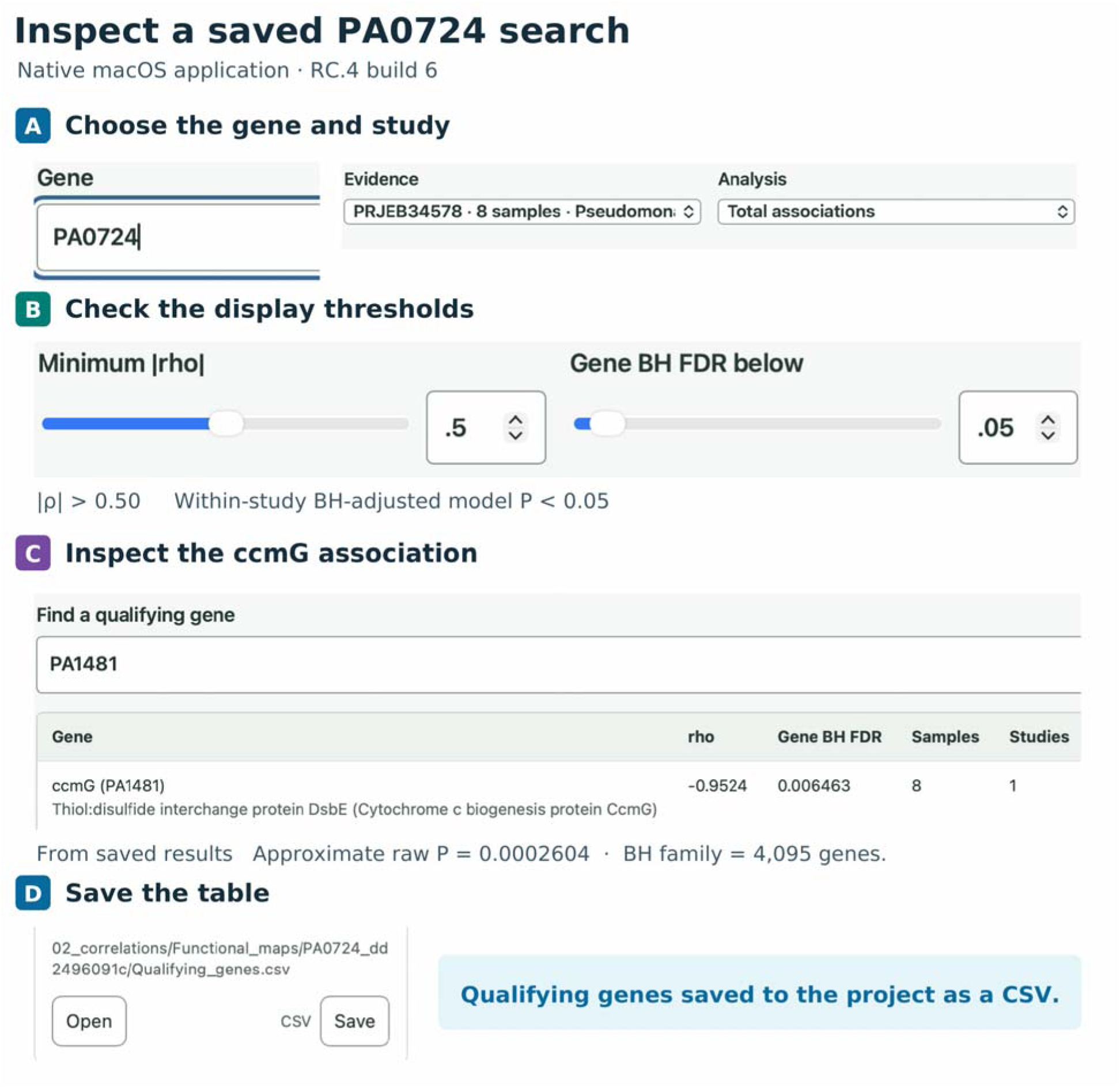
Following PA0724 and *ccmG* in PAannotate. A–D) authentic build screenshot captures of the saved PA0724 analysis for eight PRJEB34578 records, its original cutoffs, the *ccmG* result, and the project save. The approximate raw P and BH family size in C were typeset from saved output.

**Figure 5.**
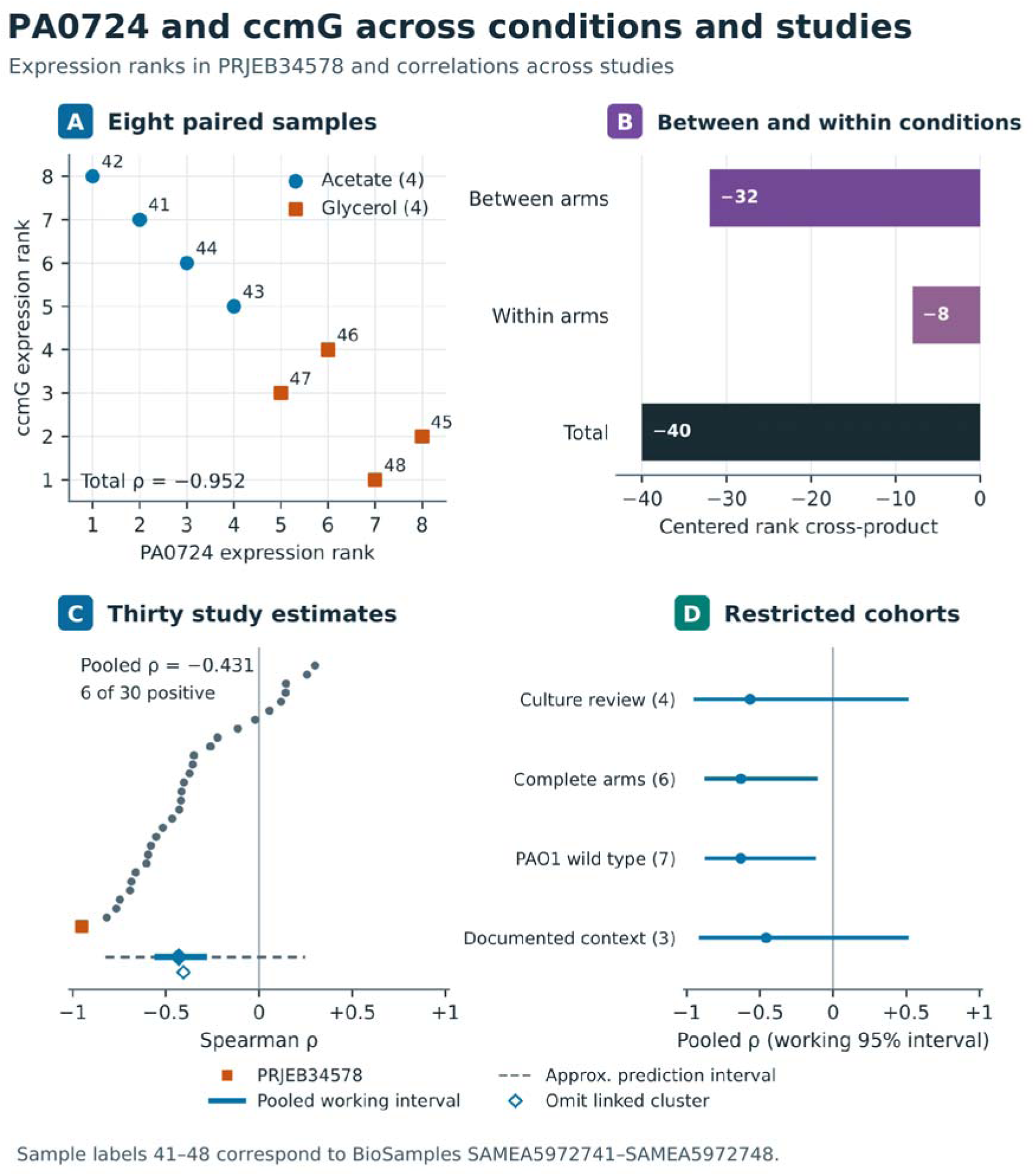
PA0724 and *ccmG* across growth conditions and studies. A) ranks for four acetate and four glycerol records from PRJEB34578. Labels 41–48 identify SAMEA5972741– SAMEA5972748. B) signed rank cross-product (−40 total, −32 between arms and −8 within arms). C) 30 study estimates (named in Fig. S2). The square marks PRJEB34578. The filled diamond gives the pooled estimate with its solid working 95% interval and dashed approximate prediction interval. The open diamond omits the cluster containing PRJEB34578. D) restricted estimates and working 95% intervals with cluster counts in parentheses.

**Figure 6.**
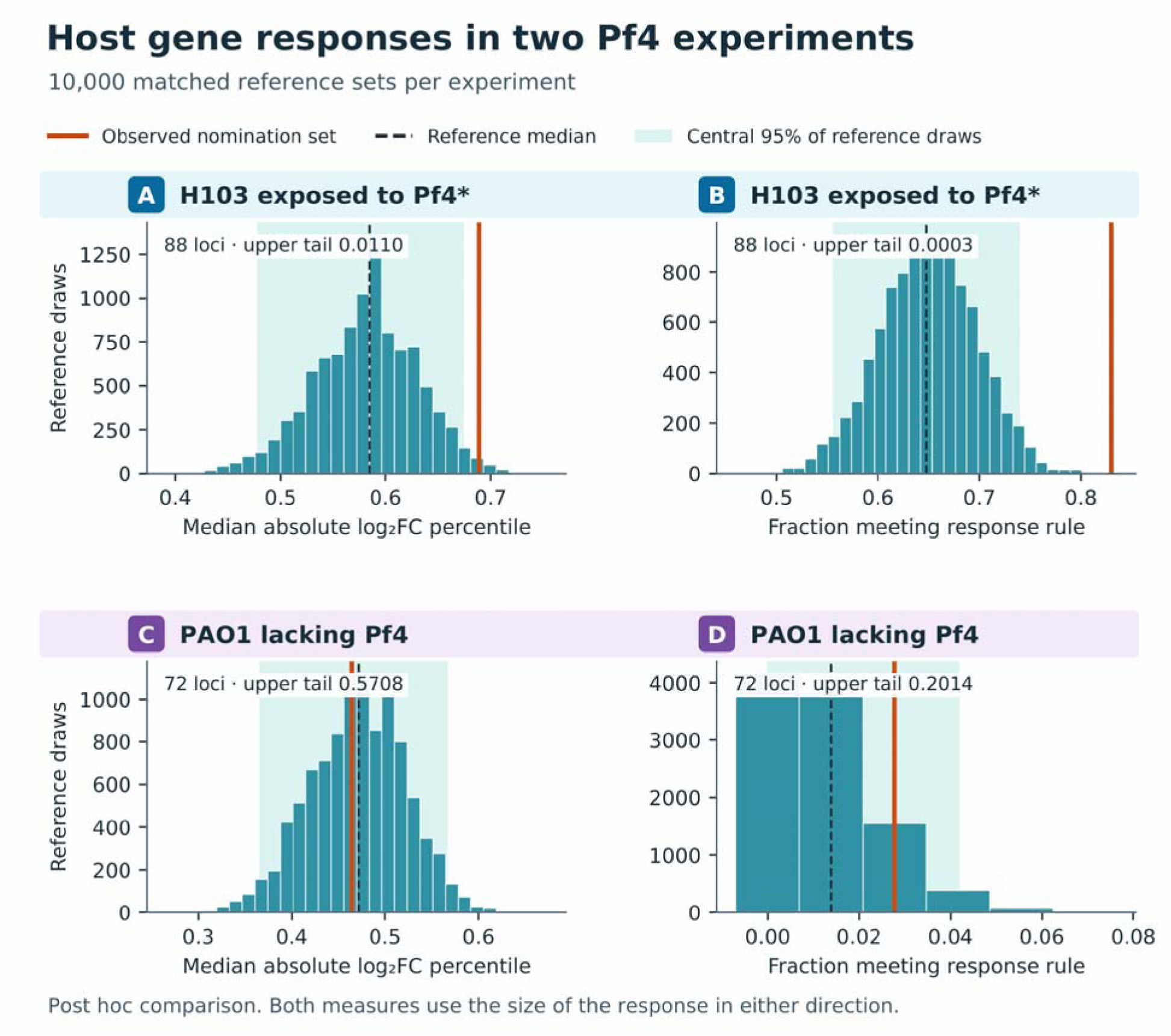
External perturbation responses and matched reference sets. A and B) H103 Pf4* exposure (88 testable nominees). C and D) PAO1 whole-Pf4 deletion (72 testable nominees). Solid lines mark the observed median absolute log fold-change percentile or response fraction. Dashed lines and shading show the median and central 95% of 10,000 post-hoc reference draws per contrast, matched on baseline detection and expression. The shading describes the reference draws. See Methods for the response rule.

## Methods

### Software implementation and local storage

The PAannotate macOS application runs the calculations on the researcher’s computer. A small program written in Swift opens the application window. It uses Apple’s AppKit for the window and WebKit to display the analysis controls within it. A service written in Python listens for requests at an address accessible only from the same computer. The service keeps a copy of the sample descriptions used for each analysis and manages the queue of requested calculations. When a researcher requests an analysis, it passes the selected samples, genes and settings to a separate R process, monitors the calculation and makes the results available to the interface. The statistical functions reside in the PAannotate R package, which can also be called directly from an R script.

The application keeps analysis records in SQLite, a database stored on disk. It saves prepared count tables and intermediate calculations as RDS files, which preserve an R data object for later use. Figure settings are stored separately from the numbers. Editing a title or changing a color can therefore redraw a saved figure without recalculating gene associations (Fig. S6). The single-study demonstration used release candidate 4 (RC.4), on an Apple silicon Mac with bundled Python 3.12.14, R 4.4.2 and PAannotate 1.0.3. We used the same bundled R environment with separate scripts for the cross-study analyses.

### From a sample search to a saved result

From the home screen (Fig. 1B), a researcher can select experiments using metadata, which are the descriptions supplied with the measurements. A search can specify the bacterial strain, the medium in which it grew, or a treatment. After inspecting the matching records, the researcher applies the selection and chooses a starting gene, called the target. A positive association means the chosen gene and another gene tend to rank high in the same samples. For a negative association, samples ranking high for one gene or signature tend to rank low for the other. In the single-study demonstration, we opened a saved search for the gene labeled PA0724 in the reference genome, inspected the gene’s statistics and the identifiers of the contributing samples, and saved the table to the project (Fig. 4). The study involved two separate growth conditions [34].

For a group of genes, the application can analyze each gene separately and show which other genes they have in common. Alternatively, a gene signature summarizes the group with one score per sample, and a user can explore what relationships that entire signature may have. To calculate the cumulative signature, the software subtracts each gene’s study mean and divides by its standard deviation, a measure of variation within that study. It takes the median of the resulting scores across the genes for each sample. Researchers can look up the functions assigned to associated genes, view relationships in a heatmap that represents their direction and strength by color, or draw a network linking genes to shared partners or functions. The eleven-gene analysis below examined each gene separately.

### Preparing sample records and comparing RNA measurements

We used the gene counts from version 0.1.0 of the Whiteley collection [8]. A read is a short sequence derived from RNA and a gene’s “count” is the number of reads assigned to it. A sequencing run is one measurement on an instrument, and the same sample can be measured more than once. Within each project we summed counts from runs assigned to the same sample, using the public BioSample identifier where available. Each resulting record refers to one sample within that project. Separate records may still come from the same original culture. For the Pf-region screen we reviewed the type of measurement, the evidence for a PAO1-related strain, documented links among samples, and whether the material came directly from a host. We set aside both external projects, including all 15 mapped records and their alternate identifiers (Supplementary Methods S1).

We put samples on a comparable scale within each project, which is a step called normalization. Samples differ in total read count, and a few abundant RNAs can take up very different shares of those reads. We used the trimmed mean of M values (TMM) method in edgeR, an R package for analyzing gene counts. TMM estimates a scaling factor for each sample from many genes after setting-aside extreme values [9, 10]. We used the scaled counts per million reads (CPM) to decide whether a gene was detected. For expression comparisons we used base-two logarithms of CPM, with edgeR’s prior count of 2 to stabilize values near zero. On a base-two logarithmic scale, a difference of one unit represents a doubling.

A project needed at least eight records for the eleven-gene screen. Each target (the genes from PA0718-PA0728 needed at least 1 CPM in half or more of the records, whereas a potential partner needed 1 CPM or more in at least one quarter of the project’s records (rounded up), with a minimum of three records. These rules avoid ranking genes that were rarely detected. We also required variation in expression, measured by the median distance of the log-CPM values from their median. The minimum was 0.05 for targets and 0.10 for partners. We excluded the genes in the range of the Pf4 prophage and identifiers with suffixes in that range from the pool of host genes to avoid finding that our targets were positively correlated with themselves. Each pair needed usable values for both genes in at least five of the same records, with variation in both genes among those records (Supplementary Methods S1–S2).

### Measuring associations within and across studies

To measure an association within a project, we used PAannotate to rank the samples by each gene’s expression, giving tied samples their average rank. Spearman’s correlation (ρ) compares the two rankings. It equals +1 if the orders are identical and −1 if one is exactly the reverse of the other. We used this measure because genes can rise together even when their amounts of RNA change by different proportions. The main analysis kept all eligible samples in a project together, including those grown under different conditions. The application obtains an approximate P value by comparing a statistic calculated from ρ with Student’s t distribution, a reference distribution determined here by the number of paired records. A two-sided P value asks how often an association at least as extreme in either direction would arise if the model held and the underlying correlation were zero.

For the saved desktop search, a study needed at least six records. The display required an absolute correlation greater than 0.50 and a Benjamini–Hochberg (BH) adjusted P value below 0.05 [13]. Absolute correlation measures strength in either direction. BH accounts for the many opportunities to find a small P value when thousands of genes are tested. Under its assumptions, it limits the expected fraction of false positives among the genes selected. We applied it to all 4,095 genes for which a correlation with PA0724 could be calculated in that study, including genes hidden by the display filter.

For the cross-study analysis, we grouped projects with documented links, such as shared samples, to reduce double counting. For each gene pair, the project with the most paired records represented its group. We combined their correlations in two steps. First, Fisher’s transformation puts correlations on a scale suitable for averaging. Second, a DerSimonian–Laird random-effects model gives greater weight to more precise estimates while allowing the underlying association to vary between studies [11]. We converted the weighted average back to the correlation scale. Supplementary Methods S2 gives the weights and the rule for correlations at or near ±1.

We used the modified Knapp–Hartung method to assess uncertainty about the mean association and to test whether that mean was zero on the transformed scale [12]. The method uses the scatter among study estimates and places a lower limit on the calculated uncertainty. The resulting 95% confidence interval describes uncertainty about the mean. We call it a working interval because its stated coverage depends on the model assumptions. When at least five study groups contributed, we also calculated a prediction interval for the underlying association in another comparable study. We adjusted the P values for all 61,259 gene pairs together using the Benjamini–Yekutieli (BY) procedure, which allows for dependence among tests [14]. We selected a pair for inspection if its absolute pooled correlation exceeded 0.30 and its adjusted P value was below 0.05. The test concerns a zero mean. Its reliability depends on the individual model P values, which we examined separately.

To see how much one group of studies influenced a result, we left out each group in turn and recalculated the pooled correlation. We also repeated the calculations for the originally selected pairs in four overlapping subsets. One retained samples traced to distinct cultures. A second retained projects in which every sample had a group label and at least two groups were represented. The other two required either source records identifying the strain as unmodified PAO1 or documentation of the medium, collection context, treatment and growth phase. We normalized the retained records again within each project and kept the original eligibility rules. Where the original records identified all experimental groups, we separated the part of the rank association attributable to differences between group means from the part arising among samples in the same group. Supplementary Methods S2 describes this calculation and the four restrictions.

### Asking which biological functions recur

We used Gene Ontology (GO), which broadly describes what individual genes do, and the Kyoto Encyclopedia of Genes and Genomes (KEGG), which groups genes into biological pathways. For each target we examined the positively and negatively associated lists separately. A one-sided hypergeometric test asks whether a list contains more genes with a given annotation than expected if the same number were drawn at random, without replacement, from the eligible annotated genes. We adjusted all 36,938 list-based tests together using BY. We also asked whether genes with a common function tended to cluster at one end of the full ranking by pooled correlation, including genes that missed the selection threshold. For this question we used fgseaMultilevel, which estimates how unusual the observed clustering is relative to random gene sets of the same size [17]. We applied BY separately to the 18,469 ranked tests (Supplementary Methods S2).

### Comparing selected genes with responses in two published experiments

We asked whether genes selected for their association with individual Pf-region genes showed unusually large changes in RNA abundance in experiments involving the whole phage. In the French study (GSE201738), Tortuel and colleagues compared strain H103 exposed to the Pf4* variant with untreated H103 [23, 62]. Ismail and colleagues compared PAO1 lacking its resident Pf4 with its parent strain (GSE154459) [24]. We used Whiteley-prepared counts for three RNA libraries per group in each comparison. A library is the material prepared from one sample for sequencing. In R, we used edgeR’s quasi-likelihood F-test for each eligible gene [9]. The test compares the difference in expression between groups with the variation among samples, allowing for uncertainty in that variation. We applied BH across the genes tested within each experiment [13]. We called a gene responsive if its expression at least doubled or halved and its adjusted P value was below 0.05.

Before examining the responses, we took up to 25 selected host genes per target and counted each gene once, yielding 88 genes. After examining the responses, we devised a comparison with 10,000 reference sets of the same testable size in each experiment. We matched reference genes to selected genes by their expression and frequency of detection in the baseline group. The change after exposure or deletion played no part in choosing the reference genes. We ranked all tested genes from zero to one by the size of their change in either direction. We then compared the median rank and the fraction of responsive genes in the selected set with those in the reference sets. The empirical upper-tail value was the number of reference sets equaling or exceeding the observed value, plus one, divided by 10,001. This comparison assesses the selected set as a whole in each experiment (Fig. 6 and Supplementary Methods S2).

### Checking calculations and statistical uncertainty

We checked numerical implementation by comparing PAannotate’s results with separately written R calculations on the same inputs. We fixed the acceptable numerical differences before comparison. Some checks used the same underlying R packages [9, 17, 18]. To check statistical calibration, we simulated data for which the underlying association was known. With a true association of zero, we counted how often an individual test gave a P value below 0.05. For intervals reported as 95%, we counted how often they contained the true mean association. These tests included an adjusted correlation that removes the estimated contribution of recorded variables before ranking the remaining expression values. The main biological screen used unadjusted correlations (Supplementary Section S5).

## Results

### Selecting experiments in the desktop application

We tested whether the desktop app could find samples from a specified experiment type and recover the selection later. A search for the Illumina HiSeq 2000 sequencing instrument returned 742 runs. Several runs could belong to one sample, so these represented 639 prepared records. Selecting strain PAO1 grown in the laboratory in LB broth--a standard bacterial growth medium--narrowed the selection to 51 records from nine projects. Reopening the saved project restored the search (Fig. 1C and Supplementary Section S4).

The same PAannotate build let us compare several target genes in two ways. When we examined the targets separately, the app showed which other genes appeared on more than one list and drew the connections between them. When we combined the targets into one expression signature, we could examine which functions were common among genes that tracked the signature. Similarly, an edited heatmap reopened with its new title and the original numerical values (Supplementary Section S4).

### How does the host respond to Pf4 exposure or deletion?

Two published experiments give a starting point for interpreting the Pf-region screen (Fig. 2D). Tortuel and colleagues found that 3,360 bacterial genes changed expression when strain H103 was exposed to Pf4*. Among the genes expressed at lower levels were those needed to make pyoverdine and pyochelin, two iron-binding compounds called siderophores [23]. Ismail and colleagues found that PAO1 biofilms with and without the resident prophage differed in their RNA profiles. In a separate measurement, biofilms lacking Pf4 produced more pyoverdine than the parent strain [24]. Iron also figures in earlier work showing that Pf4 particles can bind iron and limit its availability to the fungus *Aspergillus fumigatus* [68]. These experiments concern the whole phage. With PAannotate, we could ask a different question about its individual genes, which is whether neighboring Pf-region genes share the same expression partners.

### Do neighboring Pf-region genes share the same expression partners?

We used the R package and accompanying scripts to ask which genes elsewhere on the bacterial chromosome varied with each of the eleven Pf-region genes (PA0718–PA0728). The screen included 534 sample records from 38 projects. Documented links among those projects yielded 34 groups of potentially related studies. For each gene pair, we included at most one project from each group. Of 61,259 comparisons between a Pf-region gene and a host gene, 165 met the exploratory selection rule, involving 129 distinct host genes (Table S5).

Neighboring genes sometimes shared expression partners. PA0725 and PA0726 had 49 and 52 selected partners, respectively, with 13 in common. Figure 3A follows those 13 shared genes across all eleven Pf-region genes. To look beyond the selected lists, we compared the rankings of all 5,569 host genes common to each pair of targets. PA0722 and PA0723 ranked those genes similarly (ρ = 0.948), whereas the rankings for PA0723 and PA0728 were almost unrelated (ρ = −0.018). These correlations describe agreement between two rankings of host genes (Fig. S8).

Another shared partner, *glnS*, encodes the enzyme that attaches the amino acid glutamine to its transfer RNA (tRNA), the molecule that delivers it during protein synthesis. Both PA0725 and PA0726 were negatively associated with *glnS*. In an earlier check limited to records identified as unmodified PAO1, both estimates remained negative. Requiring a documented, distinct culture for each sample left too little evidence to determine either direction reliably (Table S9). PA0721 illustrates a different outcome. Its protein, PfsE, affects pili, the fibers on the bacterial surface, and PQS, a chemical signal that bacteria use to communicate [27, 28]. Yet none of its 5,569 host-gene comparisons met the expression-screen criteria (Fig. S4).

### Following one gene in one experiment

We returned to a single experiment to see what lay behind one of the associations in Figure 3A. This time we used PA0724 as the starting gene (Fig. 4). In the app, we reopened the saved PA0724 search for eight samples from one study (PRJEB34578). Of the 4,095 genes for which a correlation could be calculated, 1,357 met both display criteria in Methods. Of these, 722 tended to have higher expression when PA0724 expression was higher, and 635 tended to have lower expression.

One of the negatively associated genes was *ccmG* (PA1481). The CcmG protein helps keep two sulfur-bearing sites on a newly made cytochrome ready to bind heme. The iron in heme allows the mature cytochrome to carry electrons during respiration [33] (Fig. 3B). Across the eight samples, PA0724 and *ccmG* ranked in almost exactly opposite order (Spearman’s ρ = −0.9524). The approximate P value was 0.0002604 before adjustment and 0.006463 after BH adjustment across the 4,095 genes tested for PA0724 in this study. The app linked the result to the study title and eight sample identifiers (Fig. 4).

We chose this pair retrospectively from 46 associations meeting both the single-study and cross-study selection criteria. Its sample records allowed us to trace the relationship to the original growth conditions and compare it with the same pair in other studies.

### What accounted for the strong association in the eight-sample study?

The researchers grew PAO1 on acetate or glycerol as its carbon source [34]. The title named both conditions. Matching the eight sample identifiers to the original study records established which four belonged to each. All four acetate samples ranked lower for PA0724 and higher for *ccmG* than all four glycerol samples. Most of the inverse relationship across the eight samples therefore arose from the separation between the growth conditions (Fig. 5A and B). Four samples per condition fell below the minimum for estimating either relationship separately.

We then located the same eight-sample estimate in the scripted comparison across studies. Across 30 study groups and 453 paired records, the mean association remained negative but was less pronounced (ρ = −0.4315, working 95% interval −0.563 to −0.279, BY-adjusted P = 0.04259 across the 61,259 comparisons). Six of the 30 study estimates were positive. The approximate 95% prediction interval, which describes the range the model allows for another comparable study, extended from −0.826 to +0.246. Omitting the group containing the original eight samples left the pooled estimate at −0.4063 (Fig. 5C and Fig. S2). The initial study contributed to the average, but it could not account for the negative average on its own.

### How much did the choice of experiments matter?

Leaving out one study group at a time preserved the direction of all 165 selected pairs, although 12 crossed the cutoff for correlation strength. We next applied the four sample restrictions in Methods. Of the original pairs, 135 could still be estimated when we required evidence that the samples came from distinct cultures, 163 when projects had complete labels for their experimental groups, 165 when records identified the strain as unmodified PAO1, and 114 when growth and treatment conditions were more fully documented. Overall, 577 of the 660 comparisons retained at least three eligible study groups. The remaining 83 yielded no estimate (Table S11).

Restricting the samples strengthened some correlations and weakened others. Nine estimates changed sign, but each had a working 95% interval that included zero, leaving the direction uncertain. PA0724 and *ccmG* had negative estimates in all four subsets, although two of their intervals also included zero (Fig. 5D and Table S12). Even the projects with complete group labels had too few samples in any one labeled group to meet the original eight-record minimum. Each subset drew on the original records, so the four checks provide overlapping evidence.

### Which functions recur among the associated genes?

We first asked whether genes with a common function appeared unusually often among the selected partners of a Pf-region gene. None of the 36,938 list-based tests met the joint BY threshold of 0.05. A different pattern emerged when we kept every eligible gene in the ranking. Across the 18,469 ranked tests, two functions recurred for all eleven targets. Genes involved in tRNA metabolism and in attaching amino acids to tRNAs clustered toward the end of the ranking associated with lower RNA levels as Pf-region expression rose. The pattern points to genes involved in protein synthesis. Measuring how rapidly the cells made proteins would require a different experiment (Fig. S3 and Table S7).

### How did the selected genes respond in two separate experiments?

The correlations raised a different question. Would the host genes identified in the screen also show unusually large changes in RNA abundance when bacteria were exposed to phage or when a resident prophage was removed? Taking up to 25 selected genes per target yielded 88 distinct host genes. We examined them in two published experiments reserved from the association screen. Tortuel and colleagues compared strain H103 exposed to the Pf4* variant with untreated H103 [23, 62]. Ismail and colleagues compared a PAO1 strain lacking Pf4 with its parent strain [24]. All 88 genes could be tested in the exposure experiment and 72 in the deletion experiment.

We ranked genes from zero to one by the size of their expression change in either direction. In H103, the middle rank of the selected genes was 0.68908, compared with 0.58467 for genes matched on expression and detection before exposure. Only 109 of 10,000 matched sets reached the observed rank (upper-tail value 0.0110). In PAO1, the corresponding ranks were 0.46414 and 0.47160, and 5,708 of 10,000 sets reached the observed value (upper-tail value 0.5708). The response rule gave a similar contrast. After exposure, 73 of 88 genes met the rule, versus a median of 64.77% in matched sets. After deletion, 2 of 72 met it, versus a matched median of 1.39% (Fig. 6 and Tables S6 and S10).

### Checking the calculations and the uncertainty they report

All 982 comparisons against separately written R calculations agreed within the limits set before testing. The checks covered sample scaling, gene signatures, correlations, pooling across studies, adjustments for multiple tests, and analyses of sets of genes. Some used the same underlying R packages. Separate checks reproduced the adjustments for testing many genes, verified the 20,000 matched sets from the external experiments, and regenerated the original association and matched-reference plots from saved results (Supplementary Sections S5–S6).

In simulated data with 12 samples, fitting eight model terms left only four degrees of freedom, a measure of the independent information left to estimate variation. Although the true association after adjustment was zero, the adjusted Spearman test flagged 25.8% of datasets at a cutoff of 0.05 for the individual test. In a second simulation, three independent studies differed in their underlying correlations (standard deviation 0.5 on the Fisher scale). With the conventional normal approximation, which uses a bell-shaped reference curve, the DerSimonian–Laird 95% interval contained the true mean correlation in only 80.9% of datasets. The modified Knapp– Hartung procedure used in our biological screen also gave too many false positives when three simulated studies were each counted twice (Supplementary Section S5).

## Discussion

Here, we have described PAannotate, an R package and native macOS application capable of analyzing user-identified P. aeruginosa transcriptomes or for exploring the Whitely Lab database of 5,821 transcriptomes from 470 BioProjects. PAannotate lets a researcher choose both the genes to follow and the experiments in which to compare them.

To demonstrate PAannotate’s capabilities, we have explored bacterial transcriptomic signatures associated with Pf4 prophage, a pseudolysogenic phage associated with P. aeruginosa chronic skin and lung infections in people. In the Whiteley collection, the association list for PA0725 captured only thirteen of PA0726’s 52 selected partners, although the genes are neighbors. Treating the region as one expression signature would answer a different question about which genes track the group as a whole. A microbiologist studying a bacterial stress response faces the same choice and can use either approach with experiments selected for the strain or conditions of interest.

The choice of software depends on the question. ADAGE and iModulonDB are useful starting points for finding groups of genes that recur across experiments [7, 38–40, 42, 43]. COLOMBOS lets a reader follow selected genes across changes in experimental conditions, and iModulonDB also lets a reader inspect a particular gene pair under chosen conditions [4, 7, 35]. Researchers can take the Whiteley tables into R or a commercial workbench and specify their own comparisons [8, 71]. To reproduce the analysis here, they would also have to carry over the sample selections, calculate a correlation in each project and account for projects that share samples. PAannotate supplies the Whiteley-specific steps for choosing samples, calculating a relationship within each study, and returning to the records and settings behind it. In the desktop example we could reopen the result and its source records. Combining the eleven target analyses across studies required R and accompanying scripts. The desktop tests document the steps we could execute. How readily a first-time user learns them remains untested (Table S4).

A gene that varies with a Pf-region transcript across studies need not respond to every change involving Pf. The selected host genes showed unusually large changes as a group after exposure in H103. Deletion in PAO1 gave little evidence of the same pattern. The strains and interventions differed, so these comparisons cannot identify which difference accounts for the contrast. We selected the genes before examining their responses and devised the matched comparison after seeing them. The result describes these two experiments and leaves the biological role of each gene open.

This work has several limitations. A researcher can only select on details that the original studies recorded. For 418 of the 534 records in the Pf-region screen, the experimental group was unavailable in the compiled data. For 427, the original records left it unclear which samples came from separate bacterial cultures. Using one project per group of known related studies reduced double counting, but unrecorded links could remain. Gene coverage also limits the questions one can ask. PA0714.1, for example, was absent from the prepared Whiteley measurements. And because related Pf phages share sequences, assigning a short RNA read to a position in the PAO1 reference genome does not establish that Pf4 produced that RNA [45, 53].

What PAannotate preserves for a follow-up is the path from a candidate gene back to the samples that suggested it. A researcher can inspect whether a result comes chiefly from one experiment, recurs across several, or separates groups grown under different conditions. That distinction helps identify the comparison worth making next.

## Acknowledgements

P.L.B. discloses support from National Institutes of Health grant R01 HL148184-01; R01 AI12492093; R01 DC019965, K24 AI166718, P01AI1960471, and R01 EB038154 and support from the CFF and the CFRI. The contents are those of the authors and do not necessarily represent the views of the funding agencies.

## Data, Metadata, and Data Availability

PAannotate RC.4 was evaluated locally with bundled PAannotate 1.0.3 and R 4.4.2. The manuscript analyses used that R environment and additional scripts. Standalone PAannotate 1.0.4 and an internal source candidate, version 1.0.4.9001, underwent separate local checks. The candidate was exercised on synthetic data using different statistical settings from the manuscript analysis. PAannotate software and the full historical reproduction bundle are currently unavailable to readers and reviewers.

The upstream Whiteley *Pseudomonas aeruginosa* data release used in this study is version 0.1.0 [8]. The external series are GSE201738 and GSE154459. We used a pinned local preparation and versioned cohort and study-cluster definitions. Redistribution of the exact prepared inputs is pending. Reproducing the full historical analysis requires those inputs and the historical software and scripts.

