## Supplemental Methods and Tables for "PAannotate: Software for Exploring *Pseudomonas aeruginosa* Gene-Expression Associations from Transcriptomic Data"

Supplementary material for Paannotate

### Supplementary Methods S1 Which samples went into the analysis?

We selected samples from Whiteley v0.1.0 for the eleven-feature analysis PF4-PA0718-0728-v1.2-broad. Figure S1 shows the features and their coverage. Table S8 and Figure S7 give the evidence for their functions. Section S6 describes verification and access to historical inputs. Numbered references appear in the main manuscript.

#### From sequencing runs to sample records

The Whiteley v0.1.0 loader used 2-12-26_rawdataset.zip. We resolved duplicate sequencing run IDs, then matched count files to release metadata. We identified samples by BioSample accession where possible and used sample or run IDs as fallbacks. A BioSample shared across projects received a separate record in each project. We summed counts across runs for each gene within each record. This produced 5,586 features, 5,821 sample records and 470 project groups. The accession rules leave culture independence unresolved.

We excluded 111 of 5,932 records (138 technical runs) with explicitly nonpositive feature-read totals, and kept 23 records absent from the count summary. The initial manifest had 474 project groups. The 5,821 retained records covered 7,163 runs and 5,817 distinct stored identifiers: 5,815 BioSample accessions and two run-based placeholders without matching release-run metadata (RUN:SRR1538466 and RUN:SRR1538467). Three BioSamples appeared across projects, adding four project-specific records. These are full-collection counts. The upstream resource describes TPM data and provides feature counts ^[8]^. We prepared and normalized the feature counts within projects.

We selected 534 of 5,821 prepared records from 38 BioProjects and grouped the projects into 34 candidate clusters based on documented links. We reviewed the RNA assay, strain, known dependencies and evidence of direct host or ex vivo sampling. Missing nonessential metadata was allowed, and 442 records carried the limited-inclusion label. Within this group, parent-culture independence was unverified for 427 (419 accession-linked libraries and eight technical aggregates from the same experiment), including all ten named alternate-background records. The other 15 were documented earliest-trajectory baselines. We held back both external projects, all 15 mapped records and their aliases. Cluster assignments reflect documented links and leave culture independence unresolved. PA0718–PA0728 identifies eleven measured PAO1 reference features. The reads may also match homologous Pf-region sequences, so their source is uncertain for individual libraries.

We normalized read counts separately in each project using trimmed mean of M values (TMM) in edgeR::DGEList and calcNormFactors(method="TMM"), then calculated log₂ counts per million with cpm(log=TRUE, prior.count=2) ^[9,10]^. We retained unlogged counts per million (CPM) for the detection rules. Libraries with nonpositive totals fail processing. Quality checks record normalization warnings and factor fallbacks. Selecting different samples within a project can therefore change normalization and require new association estimates.

A project needed at least eight selected records, and each study estimate needed at least five pairs with finite values. To be eligible as a target, a feature needed CPM ≥1 in at least half of the project’s records and an unscaled median absolute deviation (MAD) of log-CPM ≥0.05. A host feature needed CPM ≥1 in at least max(3, ceiling(0.25n)) records and MAD ≥0.10. We excluded PA0715–PA0729 and their suffixes from the host tests. Each target score was the standardized log₂ CPM of one measured feature. This screen used single features throughout.

Missing details constrained which comparisons we could make. Medium was unknown or only curator-assigned for 394 records, RNA collection context for 380, and treatment for 330. Nine records had conflicting treatment labels. Growth phase was missing for 447 records and experimental group (arm) for 418. Nine projects had multiple recorded arms, covering 116 records. Genotype detail was incomplete for 235 PAO1 records, and Pf read attribution was unverified for all 534. Source time and temperature were blank throughout. These categories overlap.

Of 5,287 records left out, 3,174 fell outside the analysis scope: 2,560 by genetic background, 229 by challenge context, 146 by direct-host or blood sampling, 114 by mixed organisms, 110 by non-RNA assay and 15 by mixed-assay aggregation. Another 1,732 were unresolved, 366 fell below the project-size cutoff and 15 were held out for the external comparisons.

### Supplementary Methods S2 How did we compare genes and experiments?

#### Measuring associations within and across studies

Within a project, we asked whether the target and a candidate gene tended to rank high or low in the same samples. For the unadjusted analysis, we used Spearman correlation between the target score and candidate log-CPM, keeping finite pairs and giving tied values their average rank. We also checked Pearson correlation in numerical validation. The saved rho retains its exact value, even at an endpoint. For the approximate P value only, the software clips a copy to [−0.999999, 0.999999], then uses t=r√((n−2)/(1−r²)) and a two-sided Student t reference with n−2 degrees of freedom.

The adjusted analysis first fits each expression measure to the recorded experimental groups (~study_arm_id) within its project. It then ranks and correlates what remains after fitting those groups. Ranking first would produce a different statistic. Eligibility depends on the available design and residual rank. If group labels provide no usable information, the fit can reduce to an intercept alone, even when all group labels are missing. The production test still uses n−2 degrees of freedom, and base pooling uses a Fisher variance based on sample size. Section S5 shows why we treat the adjusted P and q values as exploratory.

To summarize an association across projects, we combined the study correlations on the Fisher scale. The pooling step separately clips each saved correlation to [−0.999999, 0.999999] before calculating zs=atanh(rs), where rs is the clipped value. The working variance is vs=1/(ns−3), with ns paired records in study s. Pooling uses the estimates and their variances. Study P values do not enter it. With ws=1/vs, μF=Σwszs/Σws, Q=Σws(zs−μF)² and C=Σws−Σws²/Σws. DerSimonian–Laird (DL) estimates between-study variance as τ̂²=max(0,[Q−(k−1)]/C) ^[11]^. The random-effects weights are Ws=1/(vs+τ̂²) and the pooled estimate is tanh(μ̂), where μ̂=ΣWszs/ΣWs. The conventional normal standard error is 1/√ΣWs. I² is max(0,[Q−(k−1)]/Q)×100, or zero if Q is zero. This Fisher summary of Spearman coefficients is approximate and estimates a different quantity from a correlation across all samples put together.

The main v1.2-broad analysis pooled unadjusted estimates using DL with modified Knapp–Hartung (mKH) uncertainty ^[12]^. For a sensitivity check on displayed pairs, we also used restricted maximum likelihood (REML) with mKH. REML estimates nonnegative τ² by bounded one-dimensional optimization and checks the zero boundary. For either estimator, the mKH standard error is √{max[1,ΣWs(zs−μ̂)²/(k−1)]/ΣWs}. The floor keeps it at least as large as the conventional weighted standard error. Tests and pointwise intervals use a t reference with k−1 degrees of freedom. We transform the intervals back using tanh. These analyses leave production settings unchanged. Section S5 describes what was learned about their performance under different assumptions.

For each gene pair, the eligible project with the most paired records stood for its candidate cluster. Ties were broken by project ID in lexical order. Pooling required at least three representatives. The model tests a zero mean association on the Fisher scale. We adjusted all 61,259 finite model P values together using Benjamini–Yekutieli (BY). The screen displayed pairs with |pooled rho| >0.30 and joint BY q<0.05. The magnitude cutoff applies to the observed estimate. We identified shared partners by matching exact host locus IDs. A separate check used only projects that contributed to all targets for a locus (12–31 common projects per locus), reusing the discovery records.

#### What changed when we left out a study or examined its groups?

We removed one retained cluster at a time and checked whether each displayed pair changed sign or crossed the absolute-rho cutoff. These refits did not recalculate joint BY values. Across 4,767 refits, the largest absolute change was 0.050052. We also tried a Fisher cap of ±0.99 and REML/mKH. No displayed pair had a retained study with an exact endpoint, and the alternate cap changed no displayed estimate. REML/mKH shifted pooled rho by at most 0.010130.

Where group labels were complete, we split the centered rank cross-product into contributions between and within groups (T=B+W). This shows how much of the observed rank relationship lies between groups. The calculation was possible for 943 rows across nine projects. The other 3,824 rows lacked enough information. We did not use this calculation for an adjusted test.

For PA0724–*ccmG*, we matched the eight app records to E-MTAB-8374 (PRJEB34578, ERP117503) ^[34]^. SAMEA5972741–SAMEA5972744 were acetate samples and SAMEA5972745–SAMEA5972748 were glycerol samples. We used the original library sizes and TMM factors and ranked each feature across all eight pairs. For each group, we multiplied the two group mean ranks after centering them on the overall means, weighted by group size and summed across groups. The within-group component uses deviations from group mean ranks. The two components sum to the total.

#### Checking the 165 selected pairs in narrower sets of samples

We asked how four restrictions changed the results for the original 534 records. (1) Prior culture review required a previously adjudicated culture and a unique, nonblank independent-unit key within the project. We excluded all members of duplicated keys and earliest-trajectory baselines. (2) Complete recorded arms required a group label for every record in a project and at least two distinct groups. (3) The wild-type (WT) restriction required exact source identification as PAO1 WT. (4) Documented context required source medium and treatment labels, source-supported collection context and a recorded growth phase. The subsets could overlap, and culture independence and within-group comparability remained unresolved.

We refit TMM within each selected project and kept the original eight-record minimum, target and host eligibility rules, minimum finite pair count, total Spearman method, choice of representatives and DL/mKH pooling. A pool still needed at least three eligible representatives. For k≥5, approximate prediction intervals used a t reference with k−2 degrees of freedom. No labeled group had eight records, so we did not fit associations within individual groups. These checks used the original 165 selected pairs and did not add a multiple-testing adjustment.

#### Two ways to test for shared functions with GO and KEGG

We asked two questions about function. For each signed list, did a gene set appear more often than expected in the eligible annotated background? We used a one-sided hypergeometric test and included gene sets with zero overlaps, with joint BY across 36,938 tests. Separately, did genes in a set tend to occur at one end of the full signed ranking? We ranked eligible annotated host loci by pooled Spearman rho for each of eleven targets in GO Biological Process, Molecular Function, Cellular Component and KEGG. We used fgseaMultilevel (fgsea 1.30.0), gene-set size 5–500, seed 2026110309, 1,000 initial permutations, sample size 101, serial execution and eps=10⁻¹⁰. Joint BY covered 18,469 ranked tests.

The ranked analysis reported 39 warnings that P values had reached its numerical reporting limit, one per affected annotation group. Two repeat runs reproduced the warning and one control did not. All 316 rerun term results matched the archive. All 44 archived groups contained the expected inputs and tested terms. The 351 rows at the limit kept P=10⁻¹⁰, missing log2err and joint BY q≈5.47×10⁻⁸. All 20 nominated terms (ten GO and ten KEGG) shared those values. A fixed tie-break determined their plotted order.

#### Comparing results with two experiments held out of the screen

Before comparing with the external experiments, we kept up to 25 displayed host associations per target, giving 114 rows and 88 distinct host loci. Both external BioProjects and mapped aliases were held out before screening. Tortuel (GSE201738/PRJNA832798) compared exposed with untreated H103. Ismail (GSE154459/PRJNA646382) compared whole-Pf4 deletion with wild-type PAO1. Each comparison used six mapped RNA libraries, three per group. We reserved three Ismail plasmid-repressor libraries. The plasmid-complement comparison was not completed.

Both comparisons used verified Whiteley counts from the held-out projects. The full set of features in the Tortuel workbook remained unresolved, and no verifiable author count matrix was available for Ismail. The shared processing pipeline limits independence of quantification. The baseline-first two-group fits used edgeR 4.4.0 TMM, filterByExpr, quasi-likelihood F tests and BH across tested host loci. Of the 5,569 input host loci, 5,569 were tested in Tortuel and 3,820 in Ismail. We kept signed effects and compared absolute log₂ fold-change percentiles between nominated and remaining tested genes within each contrast. The response rule required |log₂ fold-change| ≥1 and BH P <0.05.

#### Comparing nominated genes with matched reference sets

After seeing the original external results, we fixed the matching rules before drawing reference sets. Eligible controls came from the finite tested genes after removing all nominees. We matched on the number of nonzero counts across the three baseline libraries, then on distance in log₂(mean baseline CPM + 0.5) using totals for all 5,569 input host loci. We ordered candidate controls within each detection group by expression distance and locus ID. We visited nominees in order of increasing stratum slack (availability of matching controls), decreasing nearest-control distance and locus ID. At each visit we sampled uniformly from the nearest 20 available controls. Controls were unique within a draw and could recur across draws. We retained difficult-to-match nominees. No perturbation fold changes entered the matching.

In each draw we calculated the median percentile of absolute archived log₂ fold changes (average ranks for ties, divided by the number tested) and the fraction meeting the original response rule (|log₂ fold change| ≥1 and BH q<0.05). The upper tail divided one plus the number of draws at or above the observed value by one plus the number of usable draws. We retained all 10,000 draws per contrast. PCG64 seeds were 2026111101 (Tortuel) and 2026111102 (Ismail). Reference ranges used the linearly interpolated 2.5th and 97.5th percentiles. The result describes the archived contrasts. The procedure did not establish a calibrated experiment-level null.

All nominees were detected in all three baseline libraries and matching preserved detection exactly. Median absolute standardized expression differences were 0.0012 and 0.0235, and median Kolmogorov–Smirnov distances 0.0227 and 0.0278, for Tortuel and Ismail, respectively. Both passed the locked median thresholds (0.10 and 0.15). We kept Ismail PA0826.1, whose closest control differed by 0.95 on the matching scale. With three baseline libraries, expression variability was recorded for inspection and excluded from matching (Table S10).

### Supplementary Results S3 How did the findings change across samples and studies?

The eight records shown in the app also appear in the wider analysis. The saved PA0724–*ccmG* result and the PRJEB34578 study row agree exactly (rho −0.952380952380952, approximate raw P = 0.000260400024387253). The native BH-adjusted model P was 0.00646265515070182 among 4,095 tests. We chose the example retrospectively from 46 pairs meeting both selection rules. The scripted pool includes 30 representatives and 453 paired records (rho −0.431471154504928, joint BY-adjusted model P = 0.0425907820600972 among 61,259 tests). Figure S2 shows each project and its weight. In PRJEB34578, most of the negative rank cross-product was between the recorded groups (−32 of −40, with −8 within groups, Fig. 3). The internal CCMG_NUMERICAL_SOURCE_TABLES.csv identifies the archived inputs.

Across 165 selected pairs, 96 prediction intervals included zero. Removing one cluster at a time moved 12 pairs across the magnitude cutoff and changed no pooled signs. The four later metadata restrictions yielded 577 descriptive estimates, with median changes in magnitude varying by restriction (Table S11). Two of four restricted PA0724–*ccmG* intervals span zero (Table S12). Table S9 gives the earlier three-pair checks using only the prior-culture and source-identified WT rosters.

The earlier PA0718–*fimL* example used a different saved result and a widened display cutoff of 0.20. Its 25-cluster pool, source-arm ranks and screenshots remain in the historical archive. For PA0725 and PA0726, pooled rho with *glnS* was −0.44351 and −0.48817 across 23 and 25 clusters, respectively. In the prior-culture subset, the PA0725 pair could not be estimated and the PA0726 pair changed sign with wide uncertainty (Table S9). PA0721 had usable scores in 35 projects and a minimum joint BY-adjusted model P of 0.1040 across 5,569 pairs (Figs. S4 and S8, Tables S5 and S8).

The ranked tests found 2,200 target–term combinations among 18,469 with joint BY <0.05, covering 545 terms. Of these, 351 shared the reporting floor and a floor-dependent BY value of about 5.47 × 10⁻⁸. No term passed the separate selected-list test (36,938 tests, minimum BY 0.06725). Figure S3 shows the nominated terms. In the external comparisons, 73/88 exposure and 2/72 deletion nominees met the secondary response rule. Two and 2,013 matched draws reached or exceeded those fractions (upper tails 0.00030 and 0.20138, respectively, Tables S6 and S10, Fig. S9).

### Supplementary Methods and Results S4 What did we test in the macOS app?

#### How the app runs analyses and saves results

The macOS app sends analysis settings from its Swift/AppKit interface (WKWebView) to a local Python HTTP service. That service prepares data, saves metadata snapshots and manages jobs, caches and subprocesses. Rscript runs the bundled paannotate package and supplemental helpers (Fig. S6). The app saves analytical objects in SQLite and uses RDS files for prepared matrices and checkpoints. Figure sources retain identifiers, effects, P values, adjusted P values and denominators separately from display settings. This makes it possible to redraw a supported figure without fitting the model again. Supported checkpoints verify inputs and code before resuming.

We measured calculation time, elapsed time for the whole phase, and peak resident memory of child processes on an eight-core Apple M1 MacBookAir10,1 with 8 GiB RAM, macOS 26.4.1 (25E253) and two math threads. The small service test used 60 genes, 72 samples and six projects. The parser test used 5,001 genes and ten technical runs, with a catalog-dimension adapter used only for fixture integrity checks. We recorded three preparation repeats. System caches were uncontrolled, some phases overlapped, and peak child memory excludes the combined memory of concurrent processes.

On RC.3 build 4 (Apple silicon, macOS 26.4.1), we ran the single-gene explorer, separate-target comparison, combined signature and instrument-filtered explorer on real data and checked saving and reopening. RC.3 matched RC.2 on 5,585 single-target rows and 16,755 comparison records. The signature matched on selected keys, scores, 50,537 study correlations, 33,492 pooled rows, 1,356 overrepresentation and 554 ranked-enrichment rows, and 11 heatmap data sources. Importing and reimporting the RC.2 signature in RC.3 preserved its scientific objects. Tests on RC.2 also covered resuming from a checkpoint, recovering a missing table, handling a duplicate export name and editing figure appearance. In a separate UX.1 native check, the edited heatmap reopened with its new title and the underlying data were unchanged.

In RC.3, clearing a selection could leave an old metadata review available to apply. In targeted RC.4 build 6 tests, clearing the selection prevented that review from being applied. The same tests confirmed a valid write and exact undo, rejection of unsupported suggestions, separation of profiles and return of focus. RC.4 reopened the RC.3 signature with 20 of 21 stored objects unchanged. Only the container for the local record changed. The broader analytical checks above were run on RC.3.

We checked the app through scripted actions, screenshots and the controls exposed to assistive software. Some earlier views exposed incomplete information. These checks did not measure participant usability or establish keyboard-only navigation or VoiceOver usability.

Basic search matches text. In a field-specific search, a record can match any selected value within a field and must satisfy the rules across all selected fields. Rules about sequencing runs must match the same run. A matching run may belong to a sample record that combines several runs. Recorded labels can be incomplete or too limited for statistical adjustment. In RC.3, the instrument-filtered route matched 742 Illumina HiSeq 2000 runs to 639 prepared records. Adding PAO1, in vitro and LB gave 51 records in nine projects. Saving and reopening preserved the applied query.

Figure S10 shows the warning shipped with the app. Proposed changes to labels, defaults, when results are withheld and what is exported await approval and implementation.

In RC.4 build 6, we reopened the saved eight-record PA0724 result for PRJEB34578 at |rho| >0.50 and per-study BH-adjusted model P <0.05 (Fig. 4). The fit required six records per study. Searching PA1481 showed *ccmG*. The metadata preview gave the shared study title without assigning a group to each accession (Fig. S10). Saving reproduced the qualifying-gene CSV byte for byte. We did not test a download to a destination folder on this route. Pooled comparison and signature export were tested on a separately rebuilt RC.1 build 1. The 30-cluster manuscript result used scripts.

For the main native signature, calculation took 76.316 s, the phase took 80.7339 s and peak child memory was 546.9 MiB. For the frozen signature the values were 39.622 s, 43.0563 s and 294.7 MiB. Main native signature enrichment (ORA plus fgsea) took 66.629 s to calculate and 69.2264 s for the phase, with 371.8 MiB. The main supplemental phase took 374.0017 s, including 237.0447 s for DL/mKH and 93.4137 s for REML/mKH. Figure S5 also shows small-input cache and preparation tests. No competing tools were timed.

The original frozen broad native job timed out at 1,800.532 s, reaching 1,268.4 MiB peak child memory. Saved study checkpoints supported separate supplemental pooling. The app did not finish that pooled run. These measurements leave unrestricted full-collection runtime, maximum memory needs and recovery from arbitrary full-scale interruptions unresolved.

### Supplementary Methods and Results S5 Were the calculations and statistical tests reliable?

First we checked whether Paannotate agreed with reference calculations. All 982 final independent R comparisons met their tolerances: 23 for normalization or signatures, 60 for correlation, 799 for meta-analysis, 55 for multiple testing and 45 for enrichment or expression scores. A further 417 Python reference comparisons, 57 eligibility checks and 26 resampling comparisons passed. The counts are for comparisons, some of which use the same data. Run aggregation matched all 40,008 count cells in the 5,001-gene test with eight records and three projects. Tables S1–S3 summarize selected results from the internal NUMERICAL_VALIDATION_AND_CALIBRATION.csv and follow-up work.

For the v1.2-broad analysis we checked 534 keys, 38 projects, 34 candidate clusters, eleven target checkpoints and the exclusion of all 15 mapped external records. We reconstructed 402 usable project–target scores and 212,040 project-feature quality checks, and verified 1,969,780 study rows for integrity. Independent calculations checked study correlations and pooling for one deterministically selected pair per target, all primary BY values, all displayed-pair omission fits and both external edgeR families. Ranked replay was targeted (Section S2). The discovery run omitted the nested 86-record screen, broad six-record cutoff and optional combined Pf4 signature. Later nested checks covered only three pairs.

#### How we checked the calculations

Reference checks used base R count sums, direct edgeR normalization, independent signature calculations, explicit correlation and pooling formulas, and reference calls for multiple testing and enrichment. Direct package checks share edgeR, limma or fgsea with the tested route ^[9,17,18]^. Count sums had to match exactly. Typical tolerances were 10⁻¹¹ for correlation or meta-analysis, 10⁻¹² for P values, 2×10⁻⁶ for REML and 10⁻⁸ for odds ratios. Test inputs covered sparse or constant expression, missing data, ties, near-perfect or perfect correlations, insufficient observations, heterogeneous or influential studies and unavailable results.

For n=7 we enumerated 5,040 permutations with and without ties, separately from checking the arithmetic of the approximate Spearman P value. Separate cluster-resampling checks used 256 bootstrap and 256 sign-flip draws with recorded random-number settings. Before-and-after checks covered 72 effect-selection boundary cases, eight labels and four per-target effect-family fields. Regression tests, eligibility checks and numerical validation were distinct test sets.

#### How we tested false positives and interval coverage

We simulated data with known answers to see how often a test flagged a relationship when none existed under its model, and how often intervals captured the true value. The original adjusted-correlation experiment used independent normal errors, conditional Pearson correlations of zero or 0.35, and eight adjustment scenarios: no informative covariate, continuous, categorical, strong confounding, near-saturated, rank-deficient, constant and entirely missing. Designs used n=12 or 24, with ranks of 1, 2, 6 or 8. Six methods shared draws: total Pearson, total Spearman, adjusted Pearson, residualize-then-Spearman, corrected-degrees-of-freedom partial Pearson and a rank-first partial reference. The 16 settings yielded 32,000 independent datasets and 96 method/scenario rows. The corrected partial-Pearson reference remained inactive in production.

A second experiment generated independent study effects on the Fisher scale. The mean was zero or atanh(0.35), the between-project standard deviation τ was 0, 0.1 or 0.5, and k was 3, 4, 5 or 10. DL-normal, DL/mKH and REML/mKH used the same draws. The 24 settings yielded 48,000 independent datasets and 72 method/scenario rows. Both experiments used 2,000 replicates per setting and a nominal α=0.05. Together they generated 80,000 independent datasets and 336,000 method evaluations. Methods applied to the same draw are dependent. The seeds were 2026110304 and 2026110306. The separate 25-replicate pilot was excluded.

We used exact 95% binomial Monte Carlo intervals to describe uncertainty in simulated rejection and coverage rates. We flagged material inflation when rejection exceeded 0.075 and its lower bound exceeded 0.05. We flagged material undercoverage when coverage fell below 0.925 and its upper bound was below 0.95. Implied Fisher coverage in the raw-data simulation was a diagnostic and did not test reported per-study intervals.

Follow-up tests used 218,000 distinct simulated datasets for adjusted correlations, pooling from raw expression and tests that combine evidence across targets. See Tables S1–S3 for selected results and the appendix for settings, seeds and availability rules.

#### Where the tests and intervals fell short

The tested software displays exploratory statistics with a warning. In the v1.2-broad analysis, the absolute-rho cutoff filters the observed pooled estimate.

#### Adjusted correlation tests gave too many false positives

In a simulation with 12 samples and no true association after accounting for the modeled variables, adjusted Pearson flagged 30.0% of cases at a nominal 5% cutoff (95% Monte Carlo interval 28.00–32.06%). The adjusted Spearman test flagged 25.8% (23.89–27.78%). The design had rank eight, leaving four residual degrees of freedom, a setting the native eligibility rule allowed (Table S1).

With 24 samples and six groups (design rank six), the corresponding rates were 8.05% and 7.65%. With 24 samples and only an intercept, they were 5.25% and 4.50%. The experimental corrected partial-Pearson reference gave 4.70% in the near-saturated setting and agreed with 500 linear-model tests to a maximum P-value error of 3.9968×10⁻¹⁵. It remained inactive and addresses a different statistic from residual Spearman. These rates describe the tested simulations. The real cohort’s false-positive fraction remains unknown. Adjusted coefficients can describe what remains after model fitting, but a multiple-testing correction cannot repair poorly calibrated input P values.

A separate near-saturated Gaussian simulation gave 26.70% for current residual Spearman (95% Monte Carlo interval 24.77–28.70%), 9.50% when only the degrees of freedom were changed and 4.85% for corrected partial Pearson (Table S3). The earlier 25.8% comes from the original simulation and random-number stream. When error variance changed along with a continuous confounder, corrected partial Pearson flagged 21.90%, unrestricted Freedman–Lane 18.70% and HC3 6.80% (95% Monte Carlo interval 5.74–7.99%). In the 24-sample hidden-arm design, removing group labels left only an intercept and residual Spearman flagged 41.05% of conditional nulls. The missing labels left the confounder unaccounted for.

Within-group permutation and Gaussian rotation require different assumptions and target different quantities. Neither emerged as a general replacement. In the near-saturated setting, the exact permutation test could give no P value smaller than 0.125, so it had zero rejections. In a separate nearly tied test, residuals differed by 1.55×10⁻¹⁵, yet their rank correlations were −0.8904649232 and −0.9275362319. This exposed sensitivity to nearly tied ranks alongside the documented numerical matches and same-environment replay. The production rounding and tie rules stayed as implemented.

#### Conventional pooled intervals missed the true value too often

With three independent studies and a true between-study standard deviation of 0.5 on the Fisher scale, DL-normal flagged 19.1% of null cases (95% Monte Carlo interval 17.40–20.89%). Its nominal 95% intervals contained the true mean 80.9% of the time (79.11–82.60%). DL/mKH and REML/mKH gave 3.5% and 3.4% rejection, with 96.5% and 96.6% coverage, respectively. With ten studies under the same conditions, rejection and coverage were 7.65% and 92.35% for DL-normal, 4.55% and 95.45% for DL/mKH, and 4.95% and 95.05% for REML/mKH (Table S2).

The modified methods could also be conservative. With three studies and τ=0, neither flagged any of 2,000 null cases. The Fisher-scale model does not establish how either method behaves with zero-heavy expression, tied ranks or unrecorded links between projects. The primary reporting policy still requires PI and statistical review.

Both modified methods missed nominal coverage when we simulated sparse expression directly. With k=3, sample sizes 12, 30 and 48, and τ=0.5 with two possible underlying study effects, rejection and coverage were 28.05% and 71.95% for DL-normal, 9.60% and 90.40% for DL/mKH, and 9.50% and 90.50% for REML/mKH. The two modified methods’ coverage intervals had upper bounds of 91.66% and 91.75% (Table S3). In a separate three-study Gaussian test with random project signs, the derived-signature component rejected 18.55% with DL/mKH and 18.75% with REML/mKH. All nine components under each modified method met the material-inflation rule. Thus ties and zeros are only part of the problem. With five studies in the sparse design, DL/mKH and REML/mKH gave rejection rates of 7.60% and 7.85%; with ten, 4.50% and 4.55%. These simulations tested particular settings and leave the generally safe study count unresolved.

Counting each study once helped when the duplicates were known. In a Gaussian simulation that counted three unique studies as six entries, rejection was 22.85% with DL/mKH and 23.30% with REML/mKH. Representing each known cluster once reduced those rates to 2.55% and 2.60%. Unknown or partial dependence remains unresolved. In a raw Gaussian alternative with k=3, τ=0.5 and true rho=0.35, DL/mKH detected 9.90% and DL-normal detected 37.60%, with only 79.60% interval coverage for DL-normal. The results leave the choice of a primary model open.

We also asked how tests combining evidence from several targets behaved. When simulated component P values were valid, the highest rejection rate across 63 true-null settings was 5.00% (95% Monte Carlo interval 4.58–5.45%). With independent raw observations, the largest rate for the native Spearman-t conjunction was 7.05% (5.97–8.26%), in a sparse 12-sample case with independent input features and a global null (any direction, u=1). This fell below the material-inflation threshold even though its interval was entirely above nominal 5%. Direct-tail sample permutation peaked at 4.30% (3.45–5.28%). Formula checks matched 1,296 raw conjunction values to within 10⁻¹². Permutation comparisons with production were restricted to any-direction because direct directional tails differ from production halving. In the new results, 25 true-null rows had pointwise lower bounds above 5% without meeting the material-inflation rule. Twelve of these still met the upper-bound tolerance of 7.5%. These dependent summaries do not demonstrate nominal calibration.

In the three-project sign-mixture simulation, native DL-normal conjunction rejection was 24.45% at u=2. The highest mKH conjunction rates were 3.65% (DL) and 3.80% (REML) at u=9 even though their component tests were miscalibrated in the same simulation. The 199-draw permutations cannot give a P value below 0.005, well above the first-rank BY threshold of about 2.66×10⁻⁸ for 150,606 tests. These simulations establish neither biological false-discovery control nor independent replication. The Bonferroni combination can handle dependence among targets, but still needs valid component P values. The biological interpretation of the derived ninth target also remains limited.

#### Supplementary Evidence S6 What could be reproduced?

From saved results, we recalculated primary BY and external BH values, checked 20,000 matched draws and regenerated the main figures. The final figure check verified archived tables and screenshot crops. Independent replay from the exact prepared counts remains unverified. The internal archive contains exact inputs, software identifiers, source tables and verification records. Filenames here identify sources. These files do not accompany the preprint.

We installed and ran a basic local check of standalone 1.0.4. We installed the data-free reviewer candidate 1.0.4.9001 in a fresh local library under framework R 4.4.1, moved the synthetic inputs, and reopened the results in a separate R process to check five tables. The framework check reported an optional Cairo/XQuartz graphics error. A second check using an archived graphics-capable R 4.4.2 returned zero errors, zero warnings and one note for unavailable goseq. Both checks relaxed the suggested-package requirements and disabled remote incoming checks. Independent installation and replay on the original data remain unverified. Internal records: INTERNAL_INSTALLATION_AND_TEST_RECEIPT.md and REPRODUCTION_LEVELS.md.

The synthetic example has its own normalization, pooling settings and set of tests. A feature with all input counts equal to zero yielded finite correlations in three projects after preprocessing. Those correlations have no interpretable expression association. The fixture retains them for audit and excludes the feature from enrichment.

Table S1 Selected conditional null calibration cells

| **Design** | **Method** | **n / rank / residual df** | **Rejection** | **95% MC interval** |
| --- | --- | --- | --- | --- |
| **Intercept** | adjusted pearson | 24/1/23 | 0.0525 | [0.0431, 0.0632] |
| **Intercept** | adjusted spearman | 24/1/23 | 0.0450 | [0.0363, 0.0550] |
| **Intercept** | partial pearson df | 24/1/23 | 0.0525 | [0.0431, 0.0632] |
| **Six groups** | adjusted pearson | 24/6/18 | 0.0805 | [0.0689, 0.0933] |
| **Six groups** | adjusted spearman | 24/6/18 | 0.0765 | [0.0652, 0.0890] |
| **Six groups** | partial pearson df | 24/6/18 | 0.0490 | [0.0400, 0.0594] |
| **Near saturated** | adjusted pearson | 12/8/4 | 0.3000 | [0.2800, 0.3206] |
| **Near saturated** | adjusted spearman | 12/8/4 | 0.2580 | [0.2389, 0.2778] |
| **Near saturated** | partial pearson df | 12/8/4 | 0.0470 | [0.0381, 0.0572] |

Every cell contains 2,000 main-experiment replicates at nominal α=0.05 and true conditional Pearson association zero. Intercept/six-group designs use n=24; near-saturated uses n=12. The corrected partial-Pearson reference is experimental and inactive. Source: A014/F014, S0020/T020, scenario/method/true_conditional_pearson keys. Complete scenarios, alternatives, and raw implied-coverage diagnostics remain in NUMERICAL_VALIDATION_AND_CALIBRATION.csv.

Table S2 Heterogeneous Fisher working model calibration

| **k** | **Method** | **Rejection** | **95% MC interval** | **Coverage** | **95% MC interval** |
| --- | --- | --- | --- | --- | --- |
| **3** | DL normal | 0.1910 | [0.1740, 0.2089] | 0.8090 | [0.7911, 0.8260] |
| **3** | DL mKH | 0.0350 | [0.0274, 0.0440] | 0.9650 | [0.9560, 0.9726] |
| **3** | REML mKH | 0.0340 | [0.0265, 0.0429] | 0.9660 | [0.9571, 0.9735] |
| **10** | DL normal | 0.0765 | [0.0652, 0.0890] | 0.9235 | [0.9110, 0.9348] |
| **10** | DL mKH | 0.0455 | [0.0368, 0.0556] | 0.9545 | [0.9444, 0.9632] |
| **10** | REML mKH | 0.0495 | [0.0404, 0.0599] | 0.9505 | [0.9401, 0.9596] |

True mean zero and between-project Fisher-scale SD τ=0.5, not τ²=0.5; 2,000 replicates per generator cell. Rejection is at nominal 0.05; coverage is for nominal 95% intervals. Methods share generated datasets. Source: A015/F015, S0022/T022, method/k/true_tau/true_mu keys. Complete bias, width, power, and other cells remain in the numerical-validation CSV.

Table S3 Additional statistical calibration results

| **Condition** | **Method** | **Rejection** | **95% MC interval** | **Coverage** | **Result key** |
| --- | --- | --- | --- | --- | --- |
| **Near saturated n=12, rank 8** | Current residual Spearman | 0.2670 | [0.2477, 0.2870] | — | A0065 |
| **Near saturated n=12, rank 8** | Design-df residual Spearman | 0.0950 | [0.0825, 0.1087] | — | A0066 |
| **Near saturated n=12, rank 8** | Correct-df partial Pearson | 0.0485 | [0.0395, 0.0588] | — | A0067 |
| **Heteroskedastic n=24, rank 2** | Correct-df partial Pearson | 0.2190 | [0.2010, 0.2378] | — | A0083 |
| **Hidden arms n=24** | Current residual Spearman | 0.4105 | [0.3888, 0.4324] | — | A0145 |
| **Sparse k=3 n=12,30,48 τ=0.5, μ=0** | DL normal | 0.2805 | [0.2609, 0.3008] | 0.7195 | B0043 |
| **Sparse k=3 n=12,30,48 τ=0.5, μ=0** | DL mKH | 0.0960 | [0.0834, 0.1098] | 0.9040 | B0044 |
| **Sparse k=3 n=12,30,48 τ=0.5, μ=0** | REML mKH | 0.0950 | [0.0825, 0.1087] | 0.9050 | B0045 |
| **Gaussian sign mixture k=3, component 9** | DL mKH | 0.1855 | [0.1687, 0.2032] | — | C2060 |
| **Gaussian sign mixture k=3, component 9** | REML mKH | 0.1875 | [0.1706, 0.2053] | — | C2078 |
| **Ideal valid marginals PC u=3** | Bonferroni PC | 0.0500 | [0.0458, 0.0545] | — | C0012 |
| **Raw sparse IID n=12, PC u=1** | Spearman-t PC | 0.0705 | [0.0597, 0.0826] | — | C1063 |

Adjusted, raw-meta, and raw component/conjunction rows use 2,000 generated null datasets; the ideal valid-marginal row uses 10,000. Rates use evaluable tests. The hidden-arm row tests a conditional null with omitted confounding, not a marginal-null calibration claim. A/B/C abbreviate NEW_adjusted_main_/NEW_raw_meta_main_/NEW_conjunction_main_ result IDs in INFERENCE_CALIBRATION_ADDITIONAL_RESULTS.csv. Coverage is for nominal 95% intervals where defined, and dashes mean not applicable. The ideal layer tests a valid-marginal operator; the sign-mixture rows test its derived component, not conjunction. Conditions differ from the original Table S1/S2 generators. Full precision, unavailable counts, condition JSON, source hashes, and all other cells remain in the CSV. Pointwise intervals are not simultaneous; failure to meet the 0.075 material threshold does not establish nominal 0.05 calibration.

Table S4 Evidence for ten routes across eight transcriptome workflows

| **Route** | **1** | **2** | **3** | **4** | **5** | **6** | **7** | **8** |
| --- | --- | --- | --- | --- | --- | --- | --- | --- |
| **COLOMBOS ^[4,35]^** | D | D | D | R | R | D | D | D |
| **GAPE / GAUGE ^[5]^** | O | O | R | D | O | O | D | D |
| **iModulonDB ^[7,37]^** | O | O | D | R | O | O | D | D |
| **CF-Seq ^[6]^** | O | O | OR | R | O | D | O | D |
| **bacteria.guru ^[36]^** | D | R | R | R | R | D | D | D |
| **Whiteley database ^[8]^** | R | D | U | U | R | U | D | D |
| **Doing et al. ^[1]^** | R | R | D | R | R | R | D | D |
| **Paannotate desktop** | A | A | A | A | A | A | A | A |
| **paannotate R** | A | A | A | A | A | A | A | A |
| **Paannotate scripted analysis** | A | A | A | A | A | A | A | A |

Steps: 1, choose target or signature. 2, restrict by experimental metadata. 3, inspect signed expression associations. 4, combine or compare experiments. 5, inspect study contributions and possible confounding. 6, explore annotations or gene sets. 7, recover settings and data provenance. 8, export for follow-up or reproduction. D, documented. R, documented related workflow. O, observed and documented. OR, observed and documented related workflow. U, unverified. A, audited local route. The codes summarize the evidence for a particular step, including related analytical tasks. Details, versions, limits and source locators are recorded in the internal sources RELATED_TOOLS_EVIDENCE_MATRIX.csv and RELATED_TOOLS_SOURCES.md.

| **Route** | **Analytical unit and documented route** |
| --- | --- |
| **COLOMBOS** | Curated condition-contrast log ratios. Gene or module profile queries and case-specific cross-experiment partner-list intersections ^[4,35]^. |
| **GAPE / GAUGE** | Annotated GEO studies. Summarizes repeated differential expression and supports gene-set interpretation ^[5]^. |
| **iModulonDB** | Curated profiles and independent components. Gene-pair and condition-specific views, project coloring and downloadable inputs ^[7,37]^. |
| **CF-Seq** | Metadata-guided selection of a study and differential expression contrast ^[6]^. |
| **bacteria.guru** | Species-wide expression correlations and reciprocal-rank networks ^[36]^. |
| **Whiteley database** | Metadata search and retrieval of feature-count and metadata tables ^[8]^. |
| **Doing et al.** | Mapped PAO1 and PA14 compendia with documented processing and correlation analyses ^[1]^. |
| **Paannotate desktop** | Applied Whiteley selection, signed study view, saved numerical outputs, and build-specific pooled and signature routes. |
| **paannotate R** | Scripted association and pooling routes with configurable inputs and documented settings. |
| **Paannotate scripted analysis** | Frozen study clusters and weights, exact *ccmG* trace, biological restrictions and complete analysis families. |

Table S5 Target eligibility and displayed associations

| **Target** | **Projects with score** | **Positive** | **Negative** |
| --- | --- | --- | --- |
| **PA0718** | 37 | 7 | 13 |
| **PA0719** | 37 | 0 | 0 |
| **PA0720** | 37 | 0 | 0 |
| **PA0721** | 35 | 0 | 0 |
| **PA0722** | 37 | 3 | 0 |
| **PA0723** | 37 | 7 | 3 |
| **PA0724** | 36 | 4 | 5 |
| **PA0725** | 35 | 14 | 35 |
| **PA0726** | 37 | 22 | 30 |
| **PA0727** | 37 | 9 | 7 |
| **PA0728** | 37 | 5 | 1 |

Project counts refer to usable target scores among 38 selected projects. Display counts apply the joint 61,259-test BY family and the frozen effect threshold. Source: PF4_PRIMARY_AND_SENSITIVITY_RESULTS.md and PF4_ALL_ASSOCIATION_TESTS.csv.gz.

Table S6 Held-out perturbation summary

| **Contrast** | **Testable / nominated** | **Median percentile** | **Difference** | **Response** | **Tested baseline** |
| --- | --- | --- | --- | --- | --- |
| **Tortuel H103 exposure** | 88 / 88 | 0.68908 | +0.19312 | 73 / 88 | 57.30% of 5,569 |
| **Ismail PAO1 deletion** | 72 / 88 | 0.46414 | −0.03730 | 2 / 72 | 1.31% of 3,820 |

Difference is the nominated median percentile minus the median among remaining tested loci. Baseline includes all tested loci. Response required |log₂ fold-change| ≥1 and within-contrast BH P <0.05. Source: PF4_EXTERNAL_PERTURBATION_COMPARISON.csv, deduplicated_union rows.

Table S7 Selected cross-study and functional diagnostics

| **Analysis** | **Result** | **Denominator or scope** |
| --- | --- | --- |
| **Cluster omission** | 12 cross the effect cutoff, zero reverse sign | 165 displayed associations, 4,767 refits |
| **Prediction interval** | 96 include zero | 165 displayed associations |
| **Source-arm decomposition** | 943 available, 3,824 unavailable | 4,767 displayed-edge project records |
| **Selected-list GO/KEGG** | Zero with BY q<0.05 | 36,938 tests, minimum q=0.06724655 |
| **Ranked GO/KEGG** | 2,200 with BY q<0.05, 545 terms | 18,469 tests |
| **Ranked reporting floor** | 351 rows at P=10⁻¹⁰ | Common floor-dependent q≈5.47×10⁻⁸ |

Source: PF4_PRIMARY_AND_SENSITIVITY_RESULTS.md, PF4_FUNCTIONAL_ENRICHMENT_ALL_TESTS.csv and PF4_RANKED_RESULT_INTERPRETATION_HANDOFF.md.

Table S8 Evidence for the eleven reference features

| **Locus and label** | **Evidence class** | **Evidence and interpretation** |
| --- | --- | --- |
| **PA0718 PflM** | Direct phenotype | Loss of the exact locus produced loss of Pf4 lysogeny in the reported PAO1 study ^[25, Fig. 1]^. |
| **PA0719 Unassigned** | Assay result; role unassigned | A locus-level test did not abolish replicative-form synthesis in the reported context ^[29, Fig. 3 and Table 1]^. |
| **PA0720 ssDNA-binding** | Annotation / comparative | No exact-locus binding assay verified in the inspected primary set ^[29, Table 1]^. |
| **PA0721 PfsE** | Direct protein and phenotype evidence | Published experiments support interaction with PilC and effects on pilus function, and interaction with PqsA with reduced PQS signaling ^[27,28]^. |
| **PA0722 Putative pIX** | Annotation / comparative | No exact-locus functional or structural assay verified in the inspected primary set ^[29, Table 1]^. |
| **PA0723 CoaB** | Direct structure | Mature CoaB was resolved in a Pf4 filament cryo-EM structure ^[30, Fig. 1 and PDB 6TUP]^. |
| **PA0724 CoaA** | Direct interaction and phenotype evidence | Published PAO1 studies report a receptor-associated minor coat protein with partial superinfection exclusion and reported interactions with PilJ and TolR/TolA ^[27,31]^. |
| **PA0725 Putative pVI** | Annotation / comparative | No exact-locus standalone structural or functional assignment verified in the inspected primary set ^[29, Table 1]^. |
| **PA0726 Zot-like** | Annotation / comparative | pftO4-specific assembly or export was not established by a direct functional assay in the inspected evidence ^[29, Table 1]^. |
| **PA0727 RepF4** | Direct genetic evidence | Genetic evidence supports participation of cognate RepF4 in Pf4 replication ^[29, Fig. 1 and Table 1]^. |
| **PA0728 IntF4** | Direct binding and genetic evidence | PAO1 genetics and attachment-site binding support a role for IntF4 in prophage excision ^[32, Fig. 2]^. |

Source: curated locus evidence table for RefSeq NC_002516.2. Comparative Pf6 evidence ^[29]^, particle structural studies ^[30,31]^, and excision experiments ^[32]^ apply in the stated contexts. Labels identify reference features, and library-specific attribution remains unresolved.

Table S9 Historical nested checks for three selected pairs

| **Pair** | **Historical strict rho (clusters, working 95% interval)** | **Source-identified WT rho (clusters, working 95% interval)** |
| --- | --- | --- |
| **PA0718–PA1822** | −0.476 (4, −0.826 to +0.138) | −0.348 (5, −0.798 to +0.353) |
| **PA0725–PA1794** | Nonestimable (1) | −0.481 (5, −0.741 to −0.097) |
| **PA0726–PA1794** | +0.084 (3, −0.615 to +0.710) | −0.544 (6, −0.836 to −0.013) |

The strict and WT rosters contained 86 and 100 records, respectively, from seven projects each, with five and seven candidate clusters. Pair-specific eligible clusters are shown. All three WT prediction intervals included zero. These nested fixed-pair checks reused discovery data. This historical three-pair check uses two nested rosters. Tables S11–S12 describe the subsequent four-restriction analysis.

Table S10 Matched reference summary

| **Contrast and statistic** | **Observed** | **Matched median (central 95%)** | **Upper tail (exceedances)** |
| --- | --- | --- | --- |
| **Tortuel median percentile** | 0.68908 | 0.58467 (0.47890–0.67364) | 0.01100 (109) |
| **Tortuel response fraction** | 0.82955 | 0.64773 (0.55682–0.73864) | 0.00030 (2) |
| **Ismail median percentile** | 0.46414 | 0.47160 (0.36636–0.56545) | 0.57084 (5708) |
| **Ismail response fraction** | 0.02778 | 0.01389 (0.00000–0.04167) | 0.20138 (2013) |

There were 10,000 usable matched sets per contrast. Tail counts include ties. Tortuel secondary precision is limited by two exceedances. Source: PF4_EXTERNAL_MATCHED_BENCHMARK_RESULTS.csv.

Table S11 Coverage and changes under four cohort restrictions

| **Restriction** | **Estimated / 165** | **Smaller \|rho\| / median Δ\|rho\|** | **Opposite sign** | **Working interval includes 0** | **Prediction interval includes 0 / available** |
| --- | --- | --- | --- | --- | --- |
| **Prior culture adjudication** | 135 | 92 / −0.083 | 5 | 114 | 65 / 70 |
| **Complete recorded arms** | 163 | 79 / +0.009 | 0 | 94 | 112 / 133 |
| **Source-identified PAO1 WT** | 165 | 45 / +0.059 | 1 | 65 | 142 / 164 |
| **Documented context** | 114 | 68 / −0.056 | 3 | 112 | Unavailable |

Median changes compare absolute rho with the original broad estimate. Median paired-record coverage was 444 in the broad selected pairs and 64, 84, 92 and 48 in the four rows above. The 577 estimates were descriptive. The 83 nonestimable comparisons had fewer than three eligible cluster representatives. All nine opposite-sign estimates had working intervals including zero. See Table S12 for PA0724–*ccmG*. Source: SELECTED_PAIR_SENSITIVITY.csv.

Table S12 PA0724–*ccmG* under four cohort restrictions

| **Cohort restriction** | **Retained clusters** | **Paired records** | **Pooled rho** | **Working 95% interval** |
| --- | --- | --- | --- | --- |
| **Prior culture adjudication** | 4 | 54 | −0.568 | −0.953 to +0.518 |
| **Complete recorded arms** | 6 | 84 | −0.630 | −0.881 to −0.104 |
| **Source-identified PAO1 WT** | 7 | 100 | −0.631 | −0.879 to −0.116 |
| **Documented context** | 3 | 48 | −0.458 | −0.916 to +0.518 |

These archived DL/mKH fits reuse discovery records. Paired records are summed over the retained cluster representatives. Two intervals span zero. Full-precision values and cluster identifiers are recorded in the internal source table CCMG_NUMERICAL_SOURCE_TABLES.csv.

### Technical appendix for Supplementary Section S5

For reproducibility, the production REML calculation minimizes 0.5[Σ log(vs+t)+log{Σ1/(vs+t)}+Σ(zs−μ̂)²/(vs+t)] over t=τ²≥0. The first upper bound is max(1,4 var(z)). The stats::optimize tolerance is 10⁻¹⁰, with up to 12 calls and fourfold bound expansion until the optimum is below 0.95 of the upper bound. The algorithm selects zero if its objective is no larger. The independent check used upper max(10,10 var(z)) and tolerance 10⁻¹², leaving the production estimator unchanged.

#### How we generated the original simulated datasets

The adjusted-correlation simulation used seed 2026110304 and this order: none, continuous, categorical, strong, saturated, rank_deficient, constant, all_missing. Strong, saturated, constant and all_missing used n=12. The others used n=24. We drew a fresh standard-normal z before each fixed design. X contained an intercept alone (none), intercept and z (continuous or strong), an intercept with six- or eight-arm coding (categorical or saturated), intercept/z/z (rank_deficient), or intercept/zero (constant or all_missing). QR rank defined the residual dimension. The confounding term conf was zero for none/constant/all_missing, z for continuous/strong/rank_deficient, or centered dummy columns weighted 1,…,rank−1 for categorical designs. With independent standard-normal errors u,e, v=rho u+√(1−rho²)e, x=a conf+u and y=a conf+v. The multiplier a was 5 for strong and one otherwise. Rho was zero or 0.35. The shared random-number stream requires the full cell order.

For the Fisher-scale simulation (seed 2026110306), k=3,4,5,10 and study sizes were round(seq(10,60,length.out=k)). We set vs=1/(ns−3), drew zs independently from Normal(μ,vs+τ²) with μ=0 or atanh(0.35) and τ=0,0.1,0.5, then set rho=tanh(z). The separate check of native design handling used seed 2026110305.

#### How we tested the alternative methods

The follow-up generated 218,000 distinct datasets, separate from the original 80,000. Its adjusted-inference tests used ten designs with 2,000 null and 1,000 alternative draws each (30,000 datasets). Eight methods were tested: current residual Spearman, its design-df t approximation, corrected-df partial Pearson, rank-all partial correlation, unrestricted Freedman–Lane partial Pearson, within-arm midrank permutation, Gaussian residual-space rotation of the residual-rank statistic, and HC3 linear-coefficient inference. Corrected-df partial Pearson uses n−rank(X)−1 test degrees of freedom. Applying that change to residual Spearman is a separate approximation. We kept known-arm, complete-case, sparse, unequal-variance, near-saturated and hidden-arm settings separate. In the unequal-variance setting we multiplied both independent conditional errors by exp(0.8z). The complete-case test retained 18 fixed positions from 24 without changing the production missing-data rules. The original rank-first comparator used raw covariates. The follow-up ranked continuous covariates too, using a fixed standardized equally spaced covariate in place of the original normal draws. In these settings ranks are an affine transformation of the covariate, leaving other transformations untested.

Freedman–Lane shuffled residuals from the reduced model and refit nuisance effects using partial Pearson. Within-arm permutation assumes records are exchangeable within the arm. Residual-space rotation assumes spherical homoscedastic Gaussian reduced errors ^[21]^. Randomized procedures used 199 draws, P steps of 0.005 and strict P<0.05. The exact near-saturated arm test had 16 assignments and a minimum two-sided P of 0.125. A failed pilot identified an undefined HC3 calculation at leverage one, which was marked unavailable. Production methods, tie handling and eligibility were unchanged.

The pooling tests generated 60,000 raw-expression datasets across 30 settings. Methods and alternative ways of handling dependence shared draws. Gaussian population Spearman effects followed Normal(μ,τ²) on the Fisher scale, with a Gaussian-copula conversion to Pearson and sample sizes rounded along 12–48. Sparse data used 35% zeros, 20 positive levels, an integrated midrank copula mixture and two-point Fisher effects μ±τ. The effect distribution also changed, so these comparisons cannot isolate the impact of ties. Scenarios included k=3,5,10, τ=0 or 0.5, null and alternative means, n=8/k=3, and exact study duplicates counted as six ungrouped records or three known clusters. We applied variable-pair eligibility without simulating the full TMM/CPM/MAD/signature-selection pipeline.

The tests that combine evidence across targets used three kinds of inputs: 90,000 ideal vectors of valid component P values, 36,000 datasets of raw measurements and 2,000 three-project datasets. The ideal case used eight correlated Gaussian coordinates and one standardized derived coordinate, dependence 0/0.7/0.95 and zero, two or four least-favorable false nulls. The raw case used eight measurements and a fixed median-z signature without adaptive member or coherence selection. It compared native Spearman-t with IID sample permutations and direct empirical directional tails. We left uncertain derived-target boundary nulls unclassified. The three-project Gaussian stress test used n=12,24,48, primitive dependence 0.7 and symmetric project signs η=−0.7 or 0.7, with true superpopulation Fisher mean zero. The original simulations used a normal random-effects generator. Exact sign flipping across three units has eight patterns and a two-sided P floor of 0.25.

Base seeds were 2026110401 for adjusted inference, 2026110402 for raw-expression pooling and 2026110403 for conjunction tests. Reserved seed 2026110404 was unused. The internal calibration CSV indexes cell offsets and snapshots and contains 2,359 follow-up summary rows (160 adjusted, 102 raw-meta and 2,097 conjunction), with result_id, condition_json, generator and source hashes, execution receipts and counts of unavailable results. Rates use evaluable tests only. Exact pointwise binomial Monte Carlo intervals cover individual settings. The label for finite-cell noninflation uses upper bound ≤0.075 and does not establish nominal 0.05 calibration. Pilots were excluded. Whole-family biological FDR and very small P-value tails remain unvalidated.

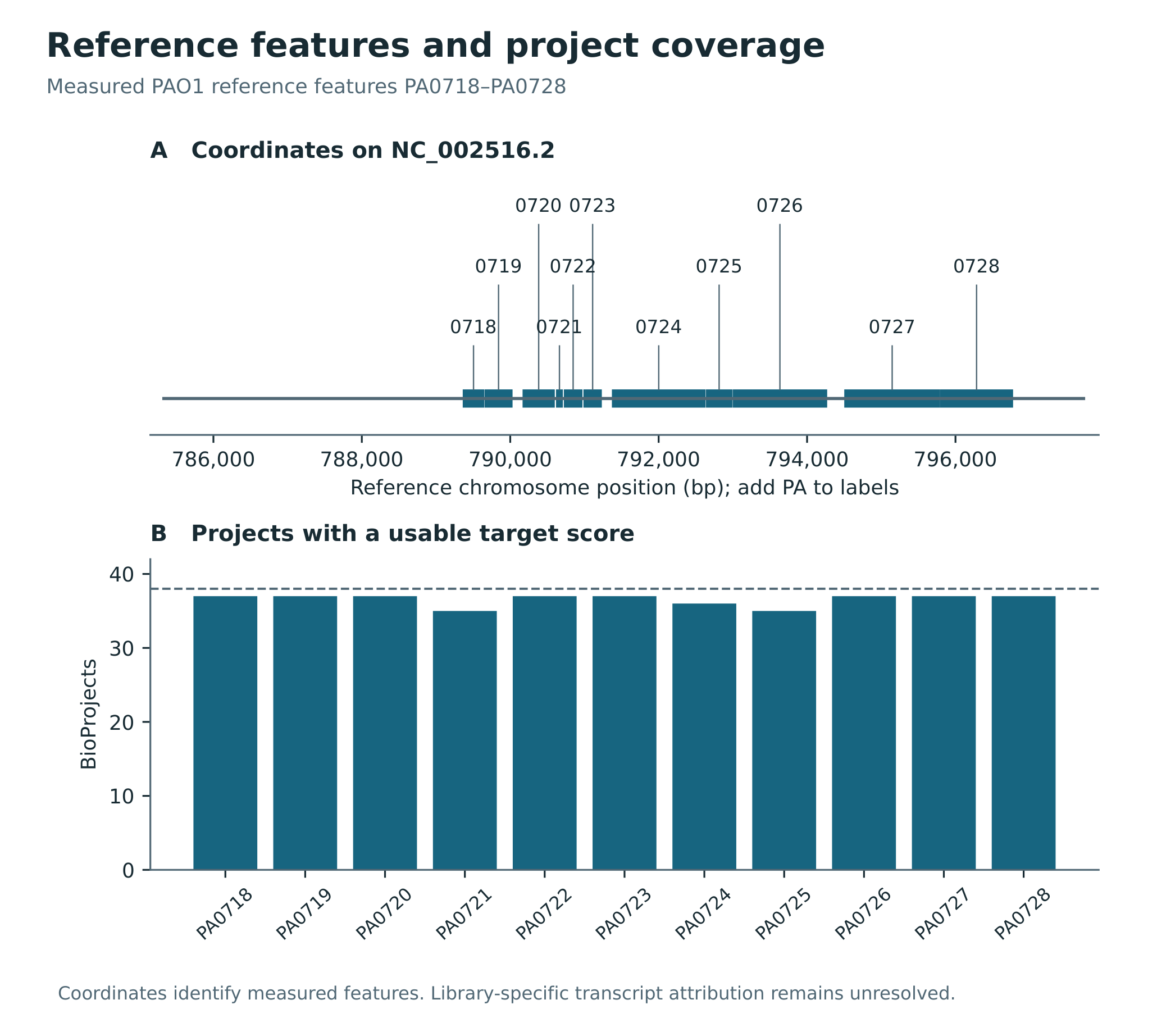

**Figure S1. Reference feature coordinates and eligibility.** A shows NC_002516.2 coordinates of all eleven measured PAO1 reference features. The horizontal line marks the previously plotted integrated reference span. B, projects with a usable target score out of 38, indicated by the dashed line. The frozen analysis retained 534 of 5,821 prepared records. Coordinates identify reference features. Library-specific read attribution remains unresolved.

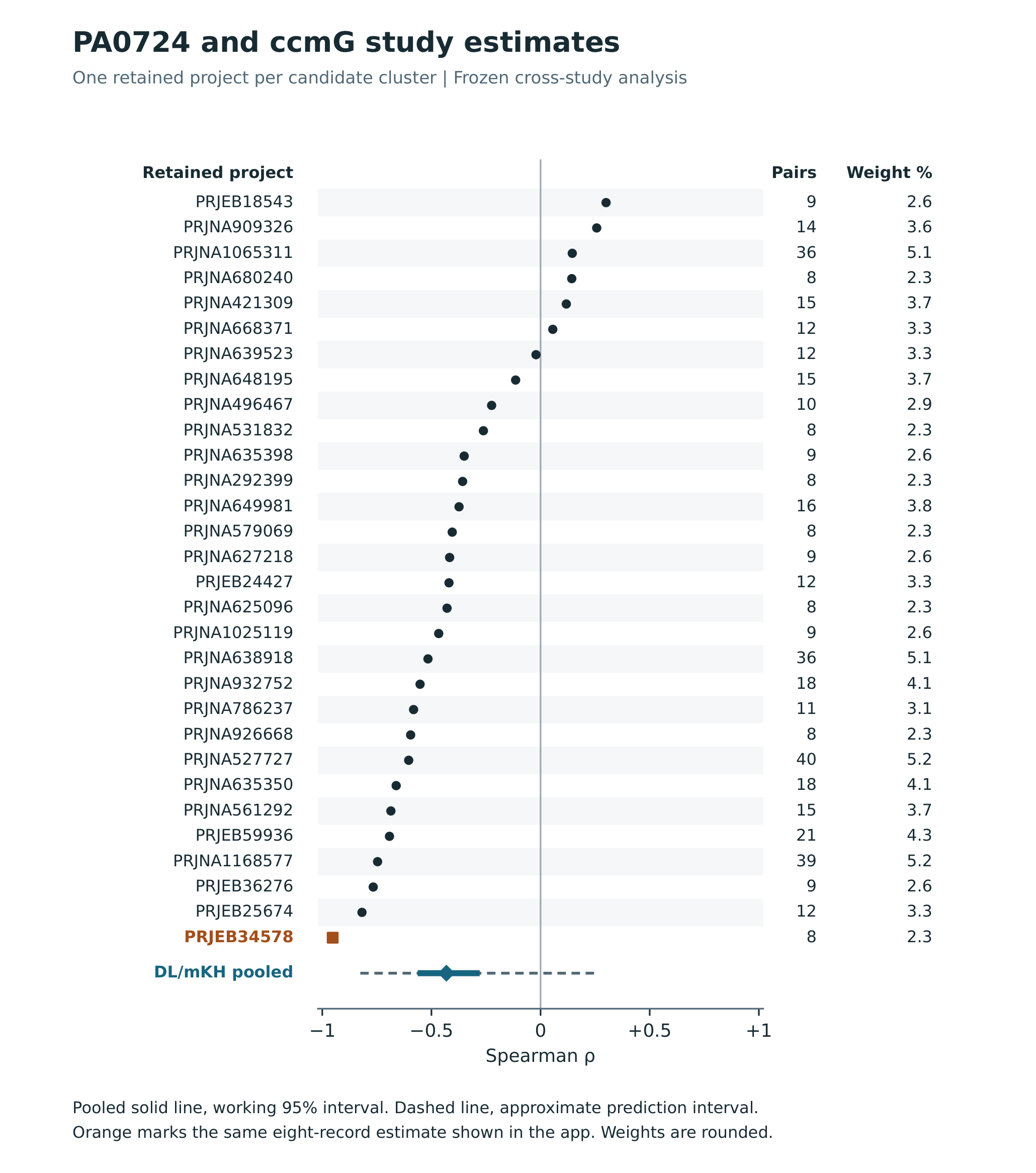

**Figure S2. Retained PA0724–*ccmG* study estimates.** One project represents each of the 30 candidate clusters. The columns give the number of paired records and each project’s share of the pooling weight. The square marks PRJEB34578. The solid and dashed segments show the pooled working and approximate prediction intervals, respectively.

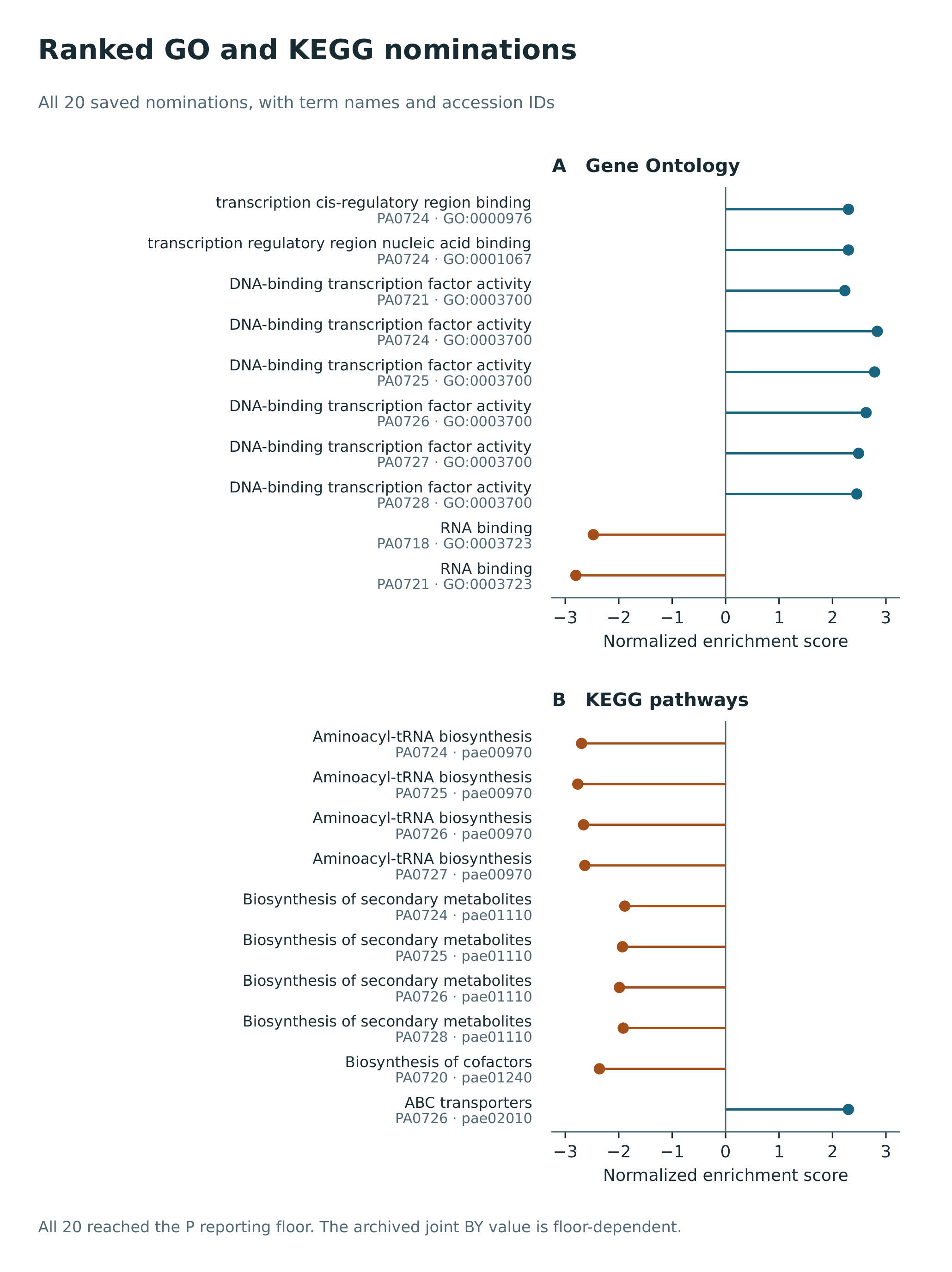

**Figure S3. Ranked GO and KEGG nominations.** Signed normalized enrichment scores show the direction and strength of enrichment for 20 previously nominated terms, ten GO and ten KEGG. All reached the P reporting floor of 10⁻¹⁰ and share a floor-dependent joint BY value. Term order follows the archived tie-break.

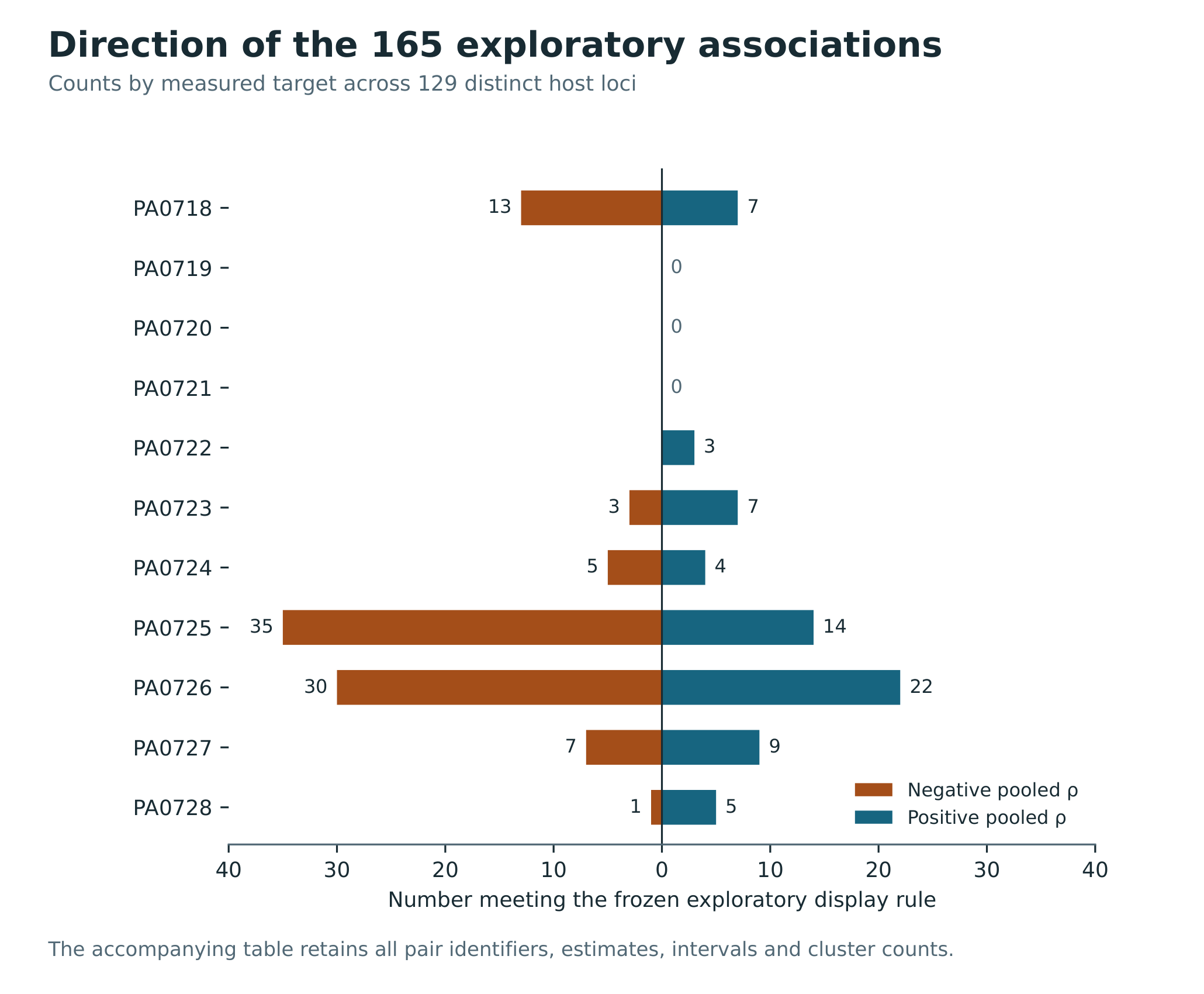

**Figure S4. Direction of the 165 selected associations.** Counts by target and sign show 165 associations across 129 host loci. The exploratory screen required |pooled rho| >0.30 and joint BY-adjusted model P <0.05. Three targets had no selected associations. These counts describe the exploratory screen.

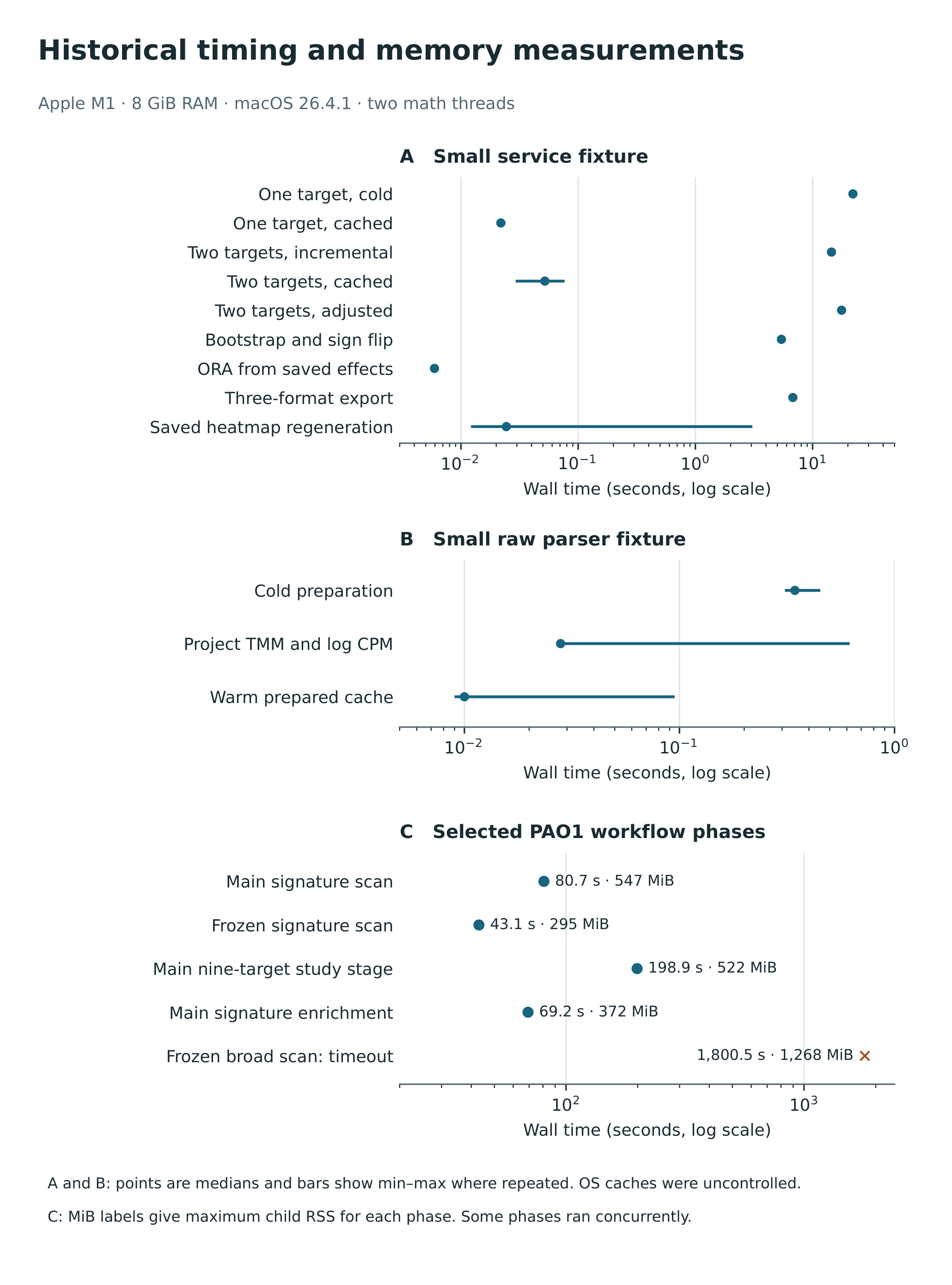

**Figure S5. Historical timing and memory measurements.** A shows a small test of the service. B shows a test of the sequencing-run parser. C shows elapsed time and peak child-process memory for selected PAO1 analysis steps, including the observed native timeout at 1,800.532 s. Time axes are logarithmic. Repeated timings show medians and ranges. See Section S4 for the measurement conditions.

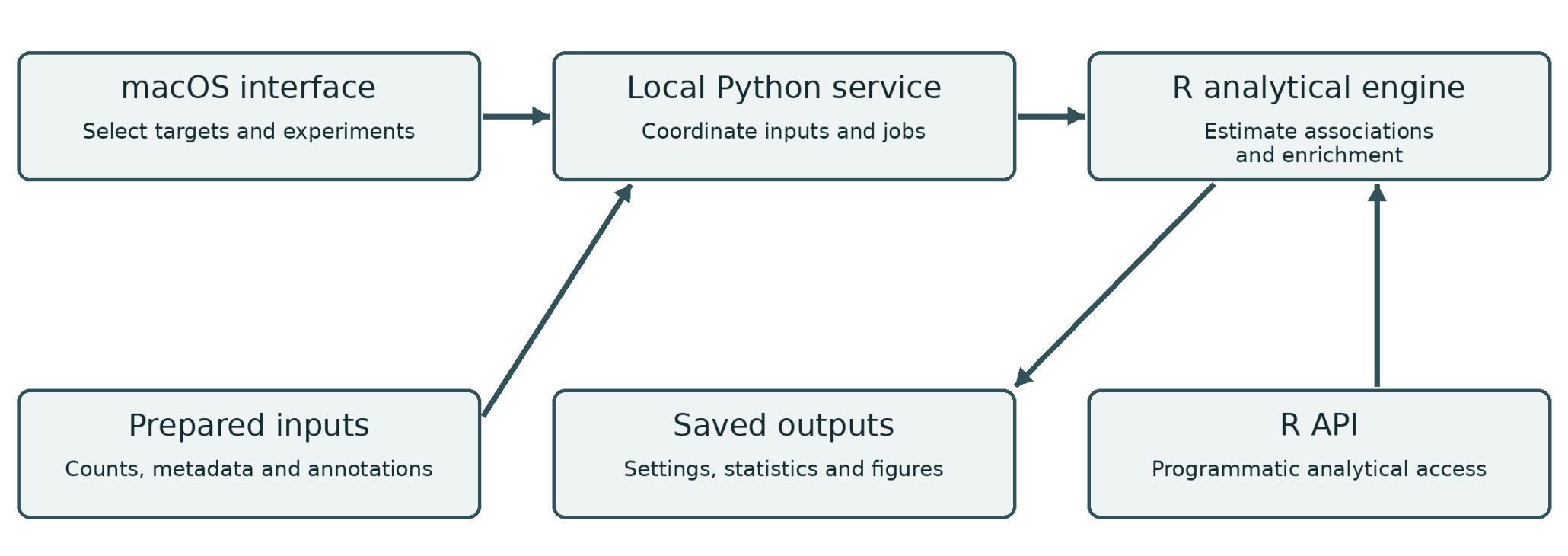

**Figure S6. Paannotate architecture.** The macOS app sends settings to a local Python service, which prepares inputs and runs the R calculations. The R package also supports scripted analyses. The diagram shows where outputs and settings are stored.

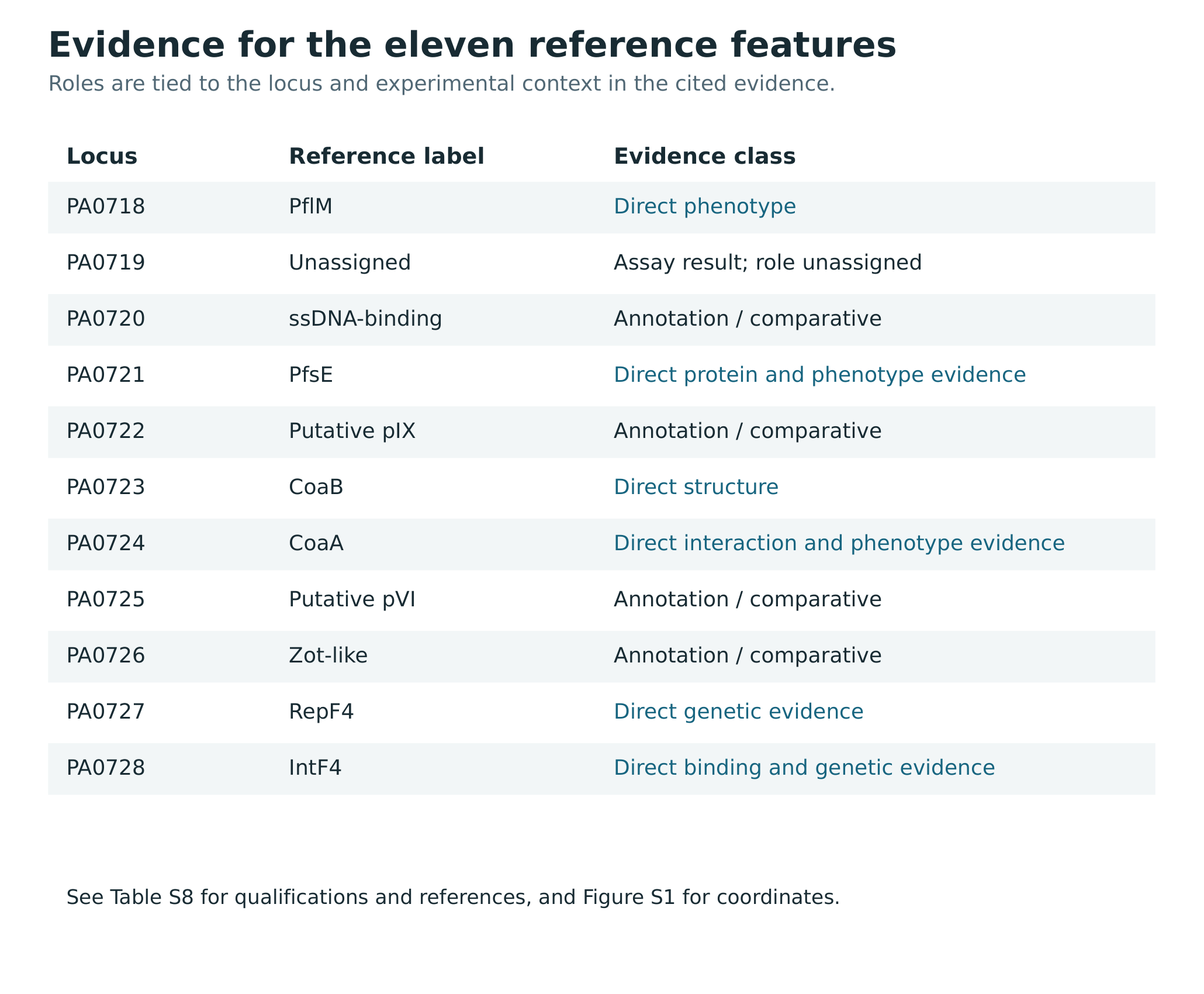

**Figure S7. Evidence for eleven reference features.** The rows give each reference feature, its short label and the type of evidence supporting it. Direct, comparative and unassigned evidence retain their original scope. See Table S8 for the full qualifications and references and Figure S1 for the reference coordinates.

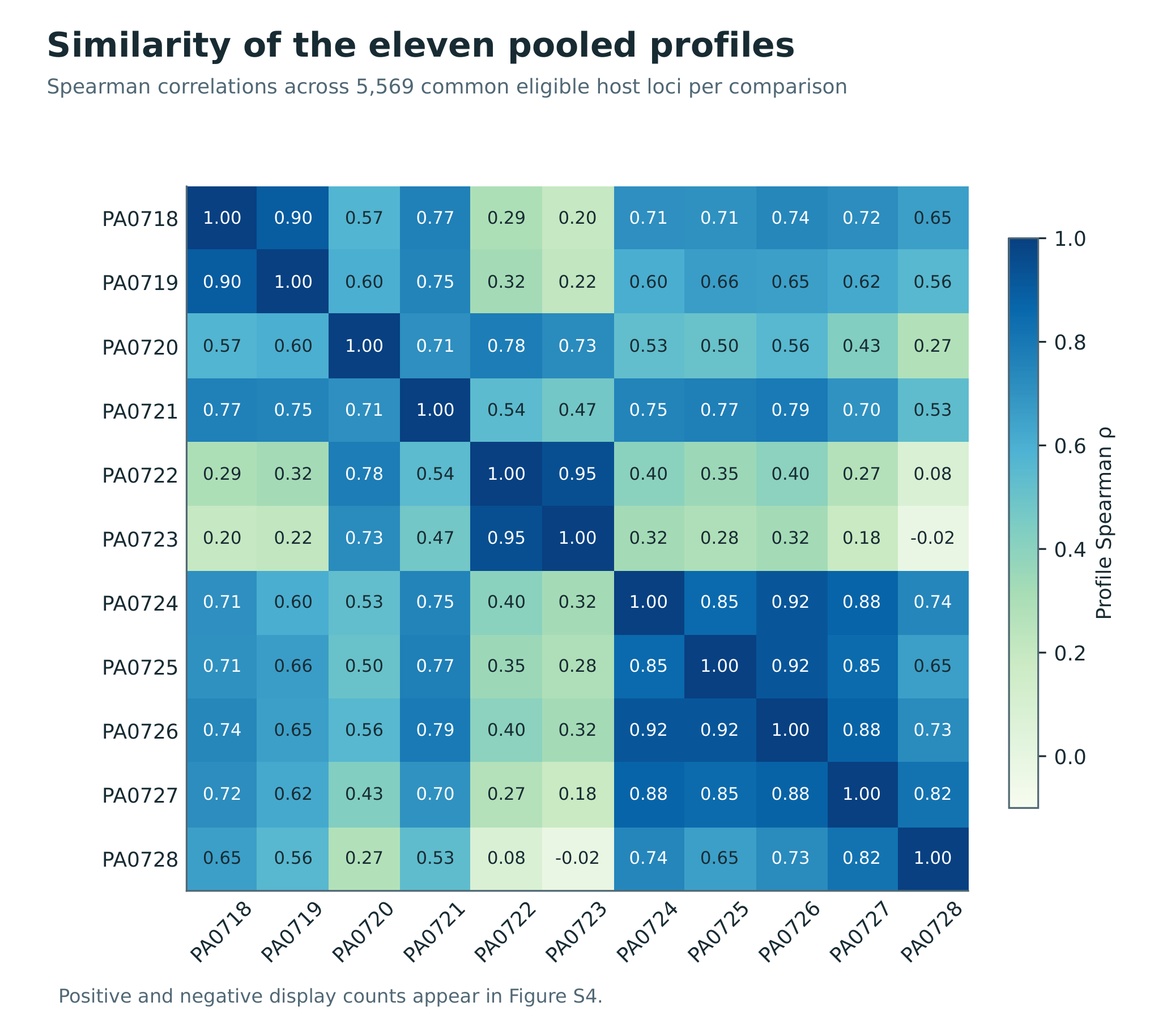

**Figure S8. Similarity of the eleven pooled profiles.** Each cell shows how closely two targets agree in their ranking of associations across 5,569 shared eligible host loci, using Spearman correlation rounded for display. Direction counts appear in Figure S4.

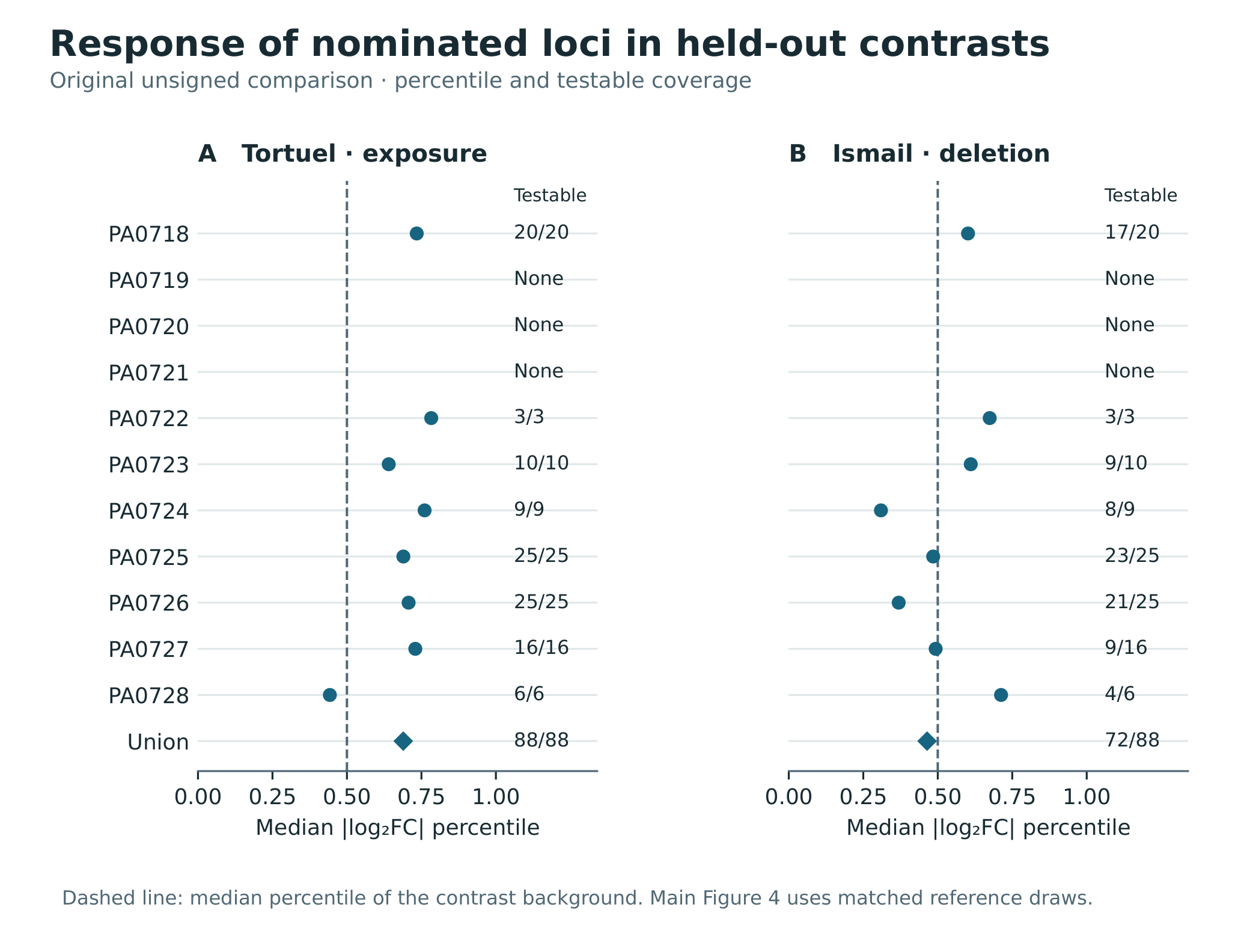

**Figure S9. Original held-out comparison by target.** The original, unmatched comparison shows the median percentile of absolute log₂ fold change among testable nominees for the Tortuel H103 exposure and Ismail PAO1 deletion contrasts. Diamonds summarize all nominated loci together, counting each locus once. Labels give testable versus nominated loci or indicate the absence of nominations. Dashed lines show the contrast background median percentile. Figure 6 uses the subsequent post hoc matched-reference benchmark.

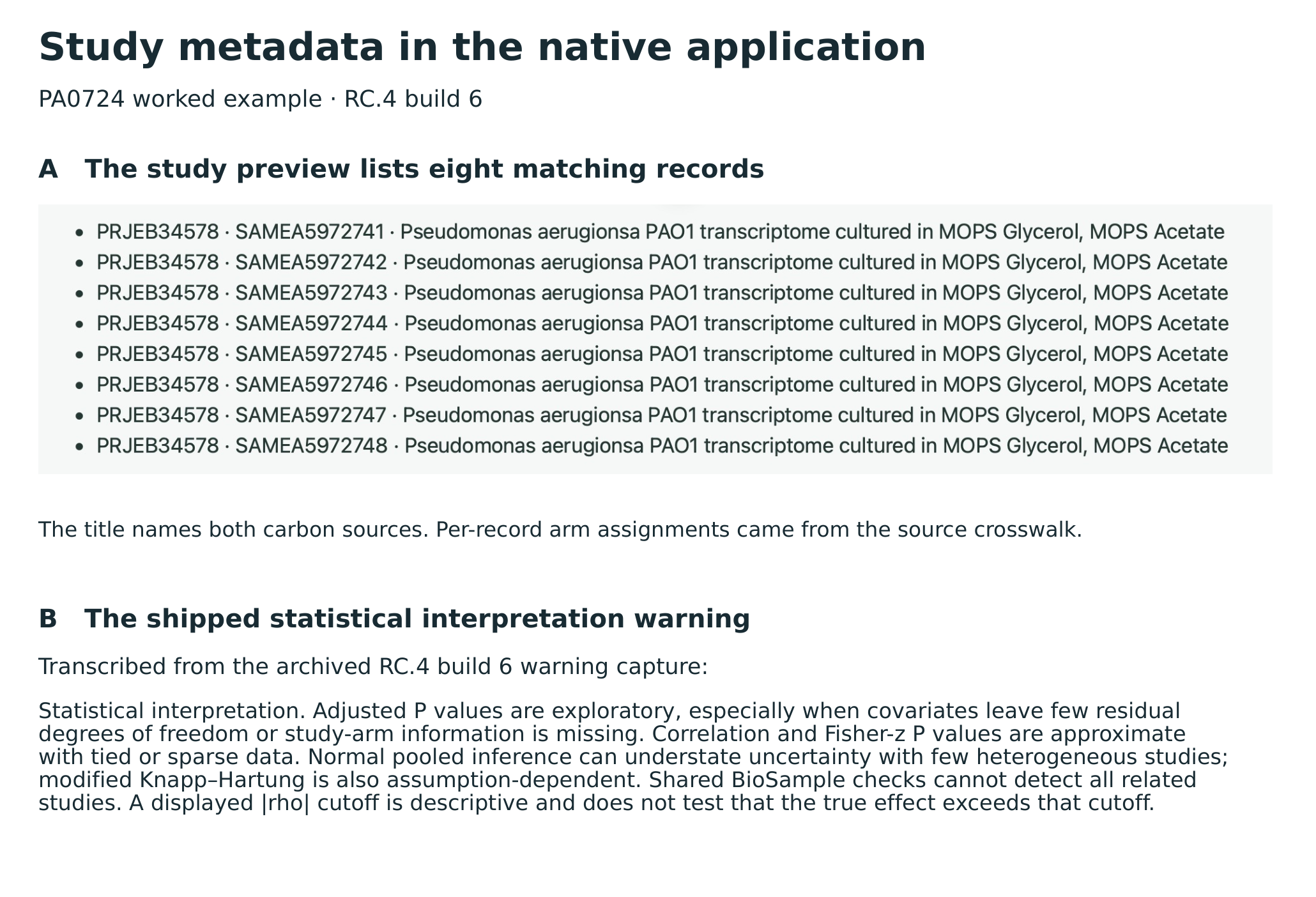

**Figure S10. Native study metadata and statistical warning.** A, eight PRJEB34578 BioSample accessions as shown in the archived PA0724 study metadata preview. The title names both carbon sources. Per-record assignments came from a match to the external study’s source records. B, the unchanged shipped statistical interpretation warning transcribed from an archived capture. The crop and transcription are identified separately.
