## Supplemental Figure S1 for "PAannotate: Software for Exploring *Pseudomonas aeruginosa* Gene-Expression Associations from Transcriptomic Data"

### Reference features and project coverage

Measured PAO1 reference features PA0718–PA0728

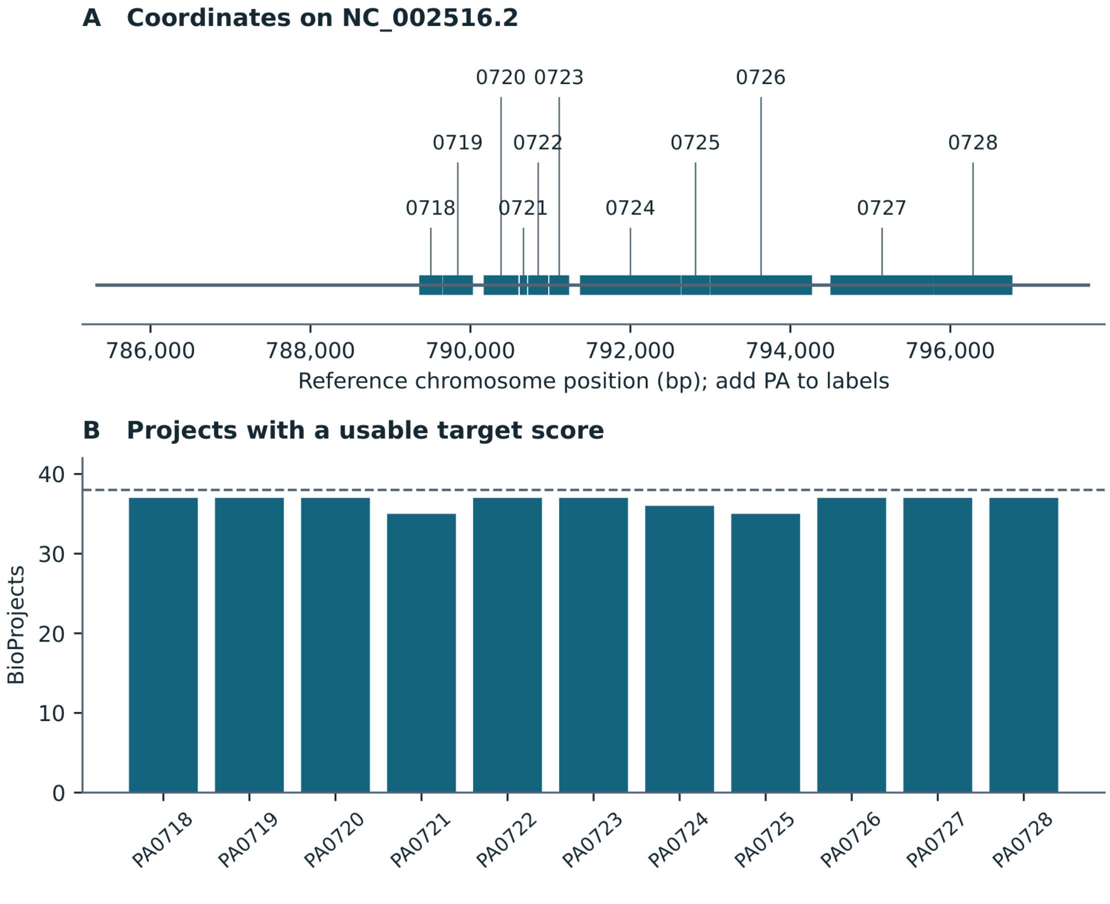


Coordinates identify measured features. Library-specific transcript attribution remains unresolved.
