## Supplemental Figure S2 for "PAannotate: Software for Exploring *Pseudomonas aeruginosa* Gene-Expression Associations from Transcriptomic Data"

### PA0724 and ccmG study estimates

One retained project per candidate cluster | Frozen cross-study analysis

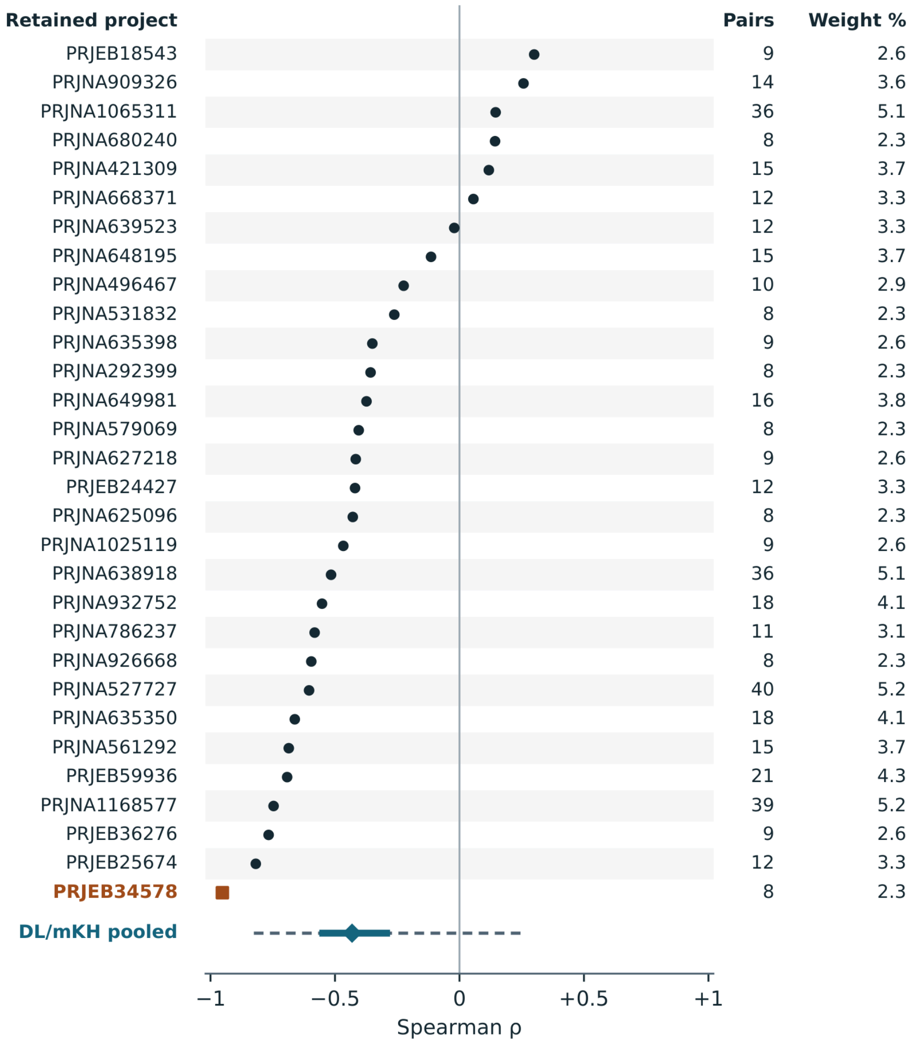


Pooled solid line, working 95% interval. Dashed line, approximate prediction interval.
Orange marks the same eight-record estimate shown in the app. Weights are rounded.
