## Supplemental Figure S3 for "PAannotate: Software for Exploring *Pseudomonas aeruginosa* Gene-Expression Associations from Transcriptomic Data"

### Ranked GO and KEGG nominations

All 20 saved nominations, with term names and accession IDs

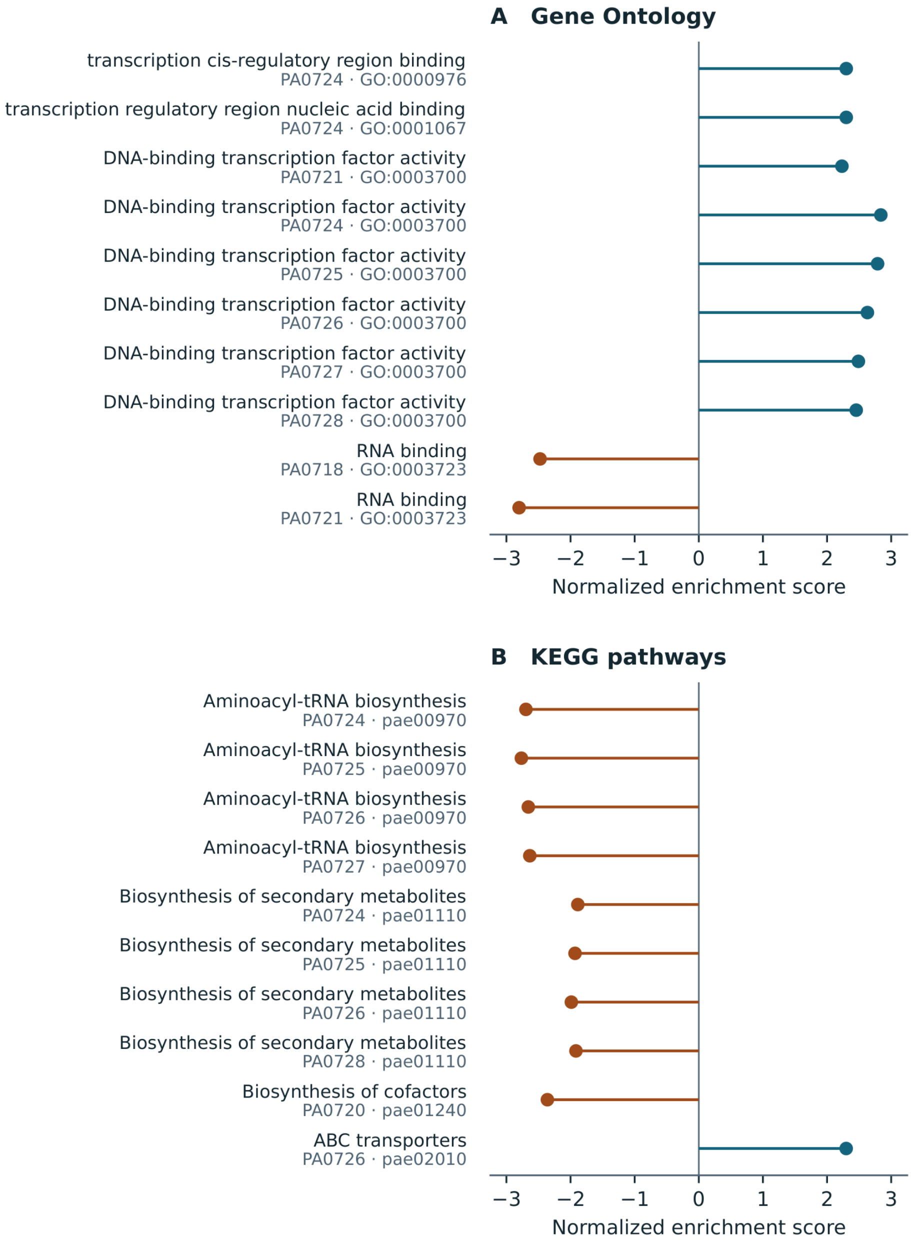


All 20 reached the P reporting floor. The archived joint BY value is floor-dependent.
