## Supplemental Figure S4 for "PAannotate: Software for Exploring *Pseudomonas aeruginosa* Gene-Expression Associations from Transcriptomic Data"

### Direction of the 165 exploratory associations

Counts by measured target across 129 distinct host loci

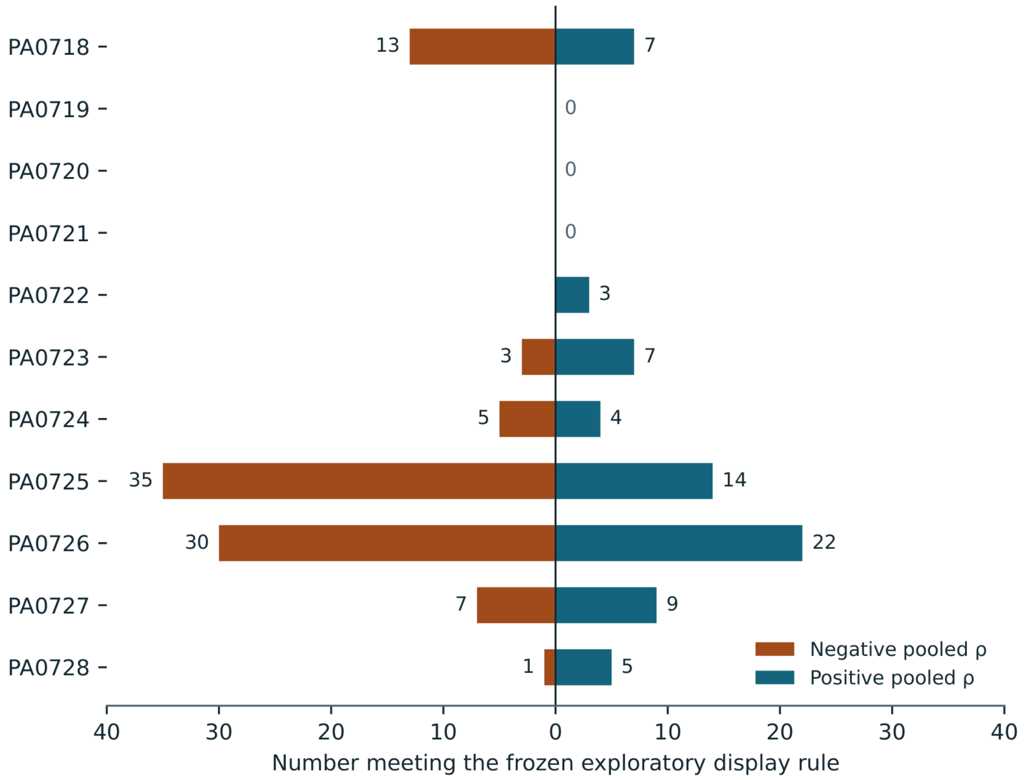


The accompanying table retains all pair identifiers, estimates, intervals and cluster counts.
