## Supplemental Figure S5 for "PAannotate: Software for Exploring *Pseudomonas aeruginosa* Gene-Expression Associations from Transcriptomic Data"

### Historical timing and memory measurements

Apple M1 · 8 GiB RAM · macOS 26.4.1 · two math threads

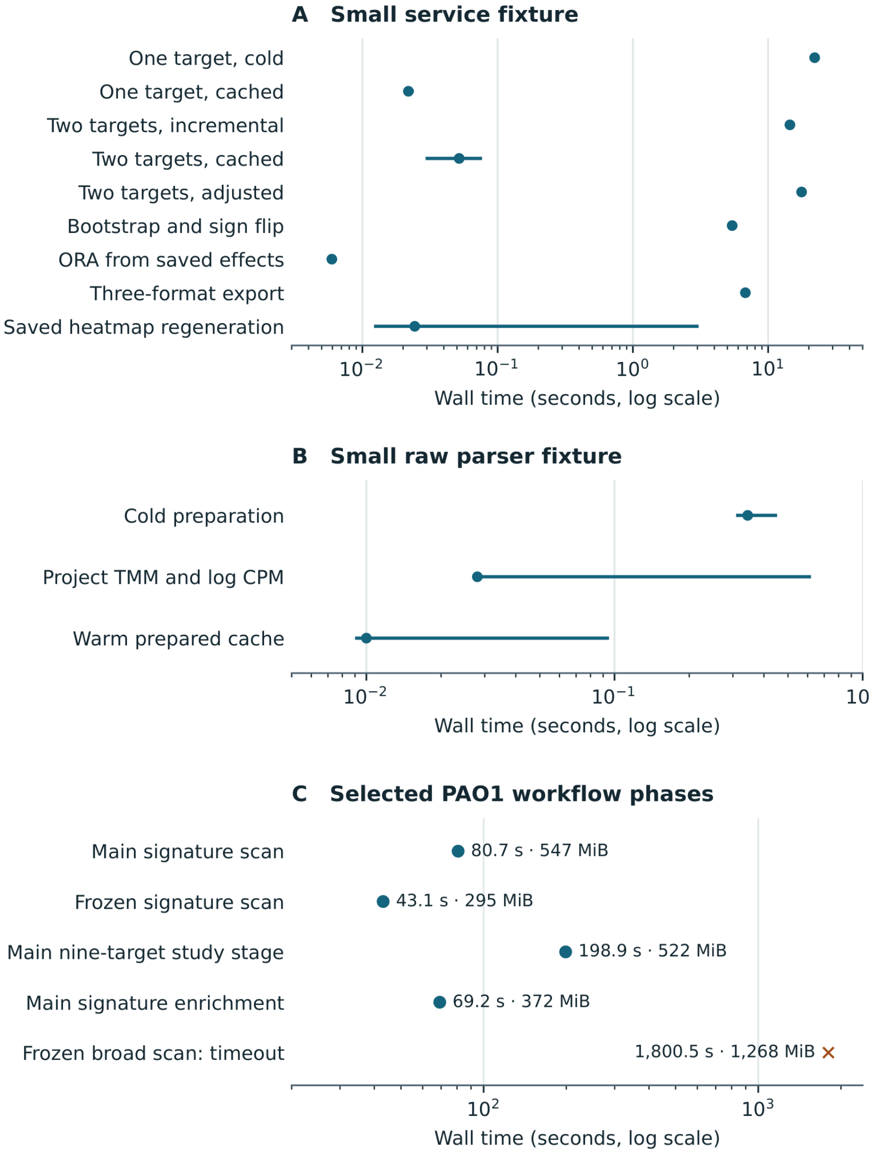


A and B: points are medians and bars show min–max where repeated. OS caches were uncontrolled.

C: MiB labels give maximum child RSS for each phase. Some phases ran concurrently.
