## Supplemental Figure S6 for "PAannotate: Software for Exploring *Pseudomonas aeruginosa* Gene-Expression Associations from Transcriptomic Data"

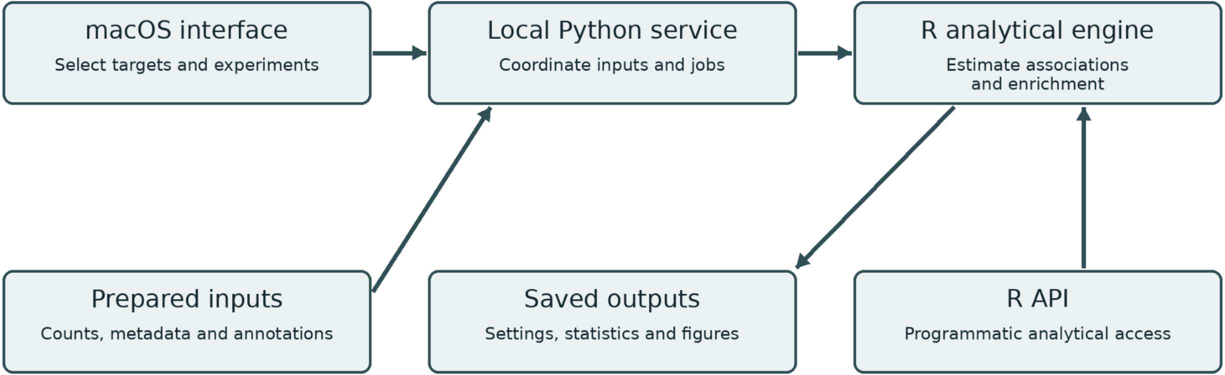
macOS interface
Select targets and experiments
Local Python service
Coordinate inputs and jobs
R analytical engine
Estimate associations and enrichment
Prepared inputs
Counts, metadata and annotations
Saved outputs
Settings, statistics and figures
R API
Programmatic analytical access
