## Supplemental Figure S7 for "PAannotate: Software for Exploring *Pseudomonas aeruginosa* Gene-Expression Associations from Transcriptomic Data"

### Evidence for the eleven reference features

Roles are tied to the locus and experimental context in the cited evidence.

| Locus | Reference label | Evidence class |
|---|---|---|
| PA0718 | PflM | Direct phenotype |
| PA0719 | Unassigned | Assay result; role unassigned |
| PA0720 | ssDNA-binding | Annotation / comparative |
| PA0721 | PfsE | Direct protein and phenotype evidence |
| PA0722 | Putative pIX | Annotation / comparative |
| PA0723 | CoaB | Direct structure |
| PA0724 | CoaA | Direct interaction and phenotype evidence |
| PA0725 | Putative pVI | Annotation / comparative |
| PA0726 | Zot-like | Annotation / comparative |
| PA0727 | RepF4 | Direct genetic evidence |
| PA0728 | IntF4 | Direct binding and genetic evidence |

See Table S8 for qualifications and references, and Figure S1 for coordinates.
