## Supplemental Figure S8 for "PAannotate: Software for Exploring *Pseudomonas aeruginosa* Gene-Expression Associations from Transcriptomic Data"

### Similarity of the eleven pooled profiles

Spearman correlations across 5,569 common eligible host loci per comparison

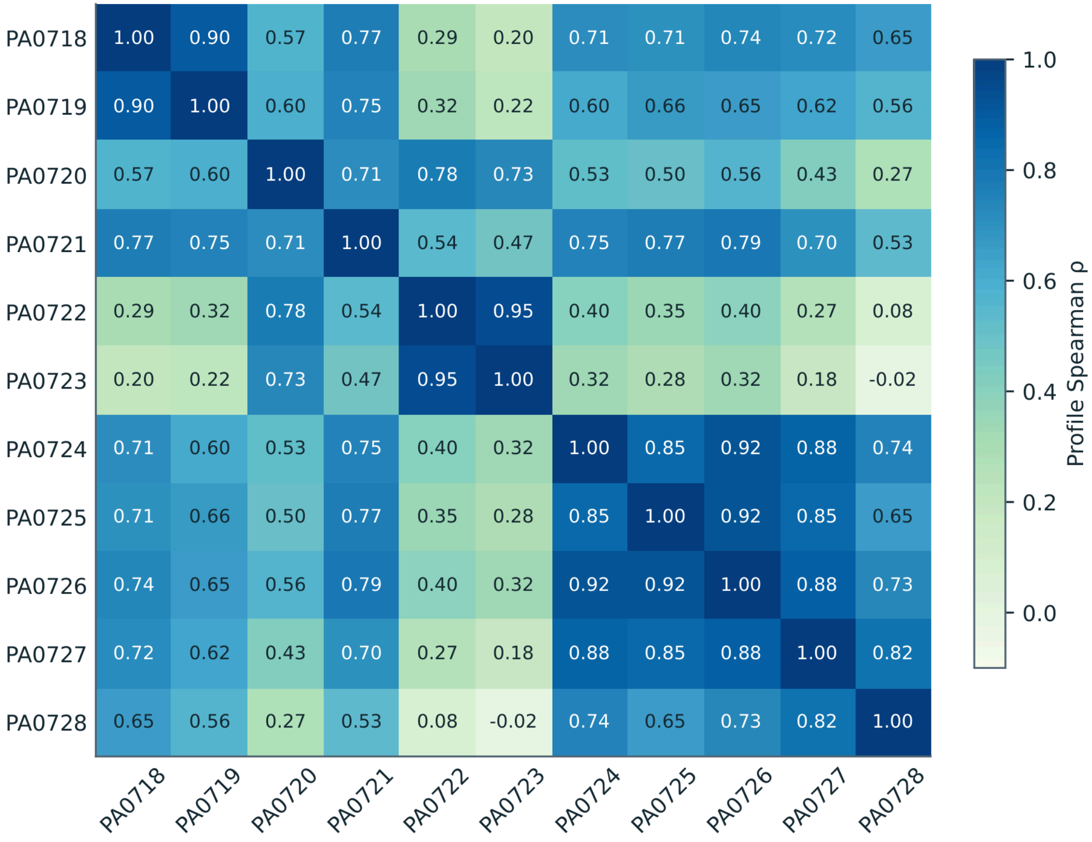


Positive and negative display counts appear in Figure S4.
