## Supplemental Figure S9 for "PAannotate: Software for Exploring *Pseudomonas aeruginosa* Gene-Expression Associations from Transcriptomic Data"

### Response of nominated loci in held-out contrasts

Original unsigned comparison · percentile and testable coverage

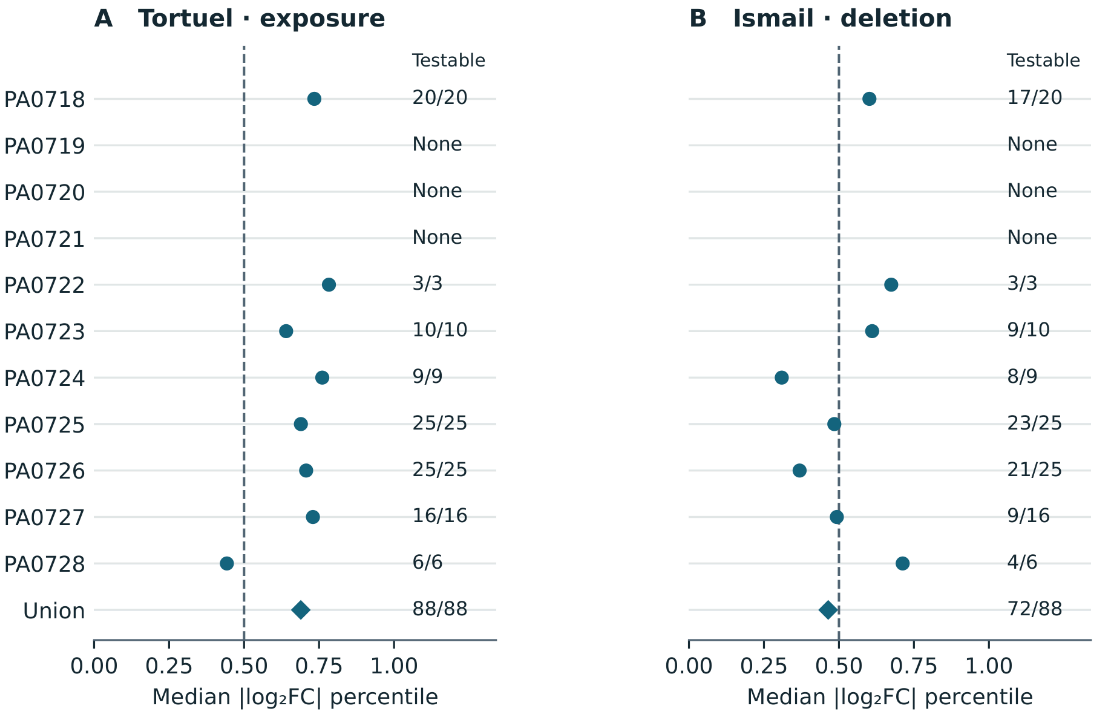


Dashed line: median percentile of the contrast background. Main Figure 4 uses matched reference draws.
