## Supplemental Figure S10 for "PAannotate: Software for Exploring *Pseudomonas aeruginosa* Gene-Expression Associations from Transcriptomic Data"

### Study metadata in the native application

PA0724 worked example · RC.4 build 6

#### A The study preview lists eight matching records

- PRJEB34578 · SAMEA5972741 · Pseudomonas aerugionsa PAO1 transcriptome cultured in MOPS Glycerol, MOPS Acetate
- PRJEB34578 · SAMEA5972742 · Pseudomonas aerugionsa PAO1 transcriptome cultured in MOPS Glycerol, MOPS Acetate
- PRJEB34578 · SAMEA5972743 · Pseudomonas aerugionsa PAO1 transcriptome cultured in MOPS Glycerol, MOPS Acetate
- PRJEB34578 · SAMEA5972744 · Pseudomonas aerugionsa PAO1 transcriptome cultured in MOPS Glycerol, MOPS Acetate
- PRJEB34578 · SAMEA5972745 · Pseudomonas aerugionsa PAO1 transcriptome cultured in MOPS Glycerol, MOPS Acetate
- PRJEB34578 · SAMEA5972746 · Pseudomonas aerugionsa PAO1 transcriptome cultured in MOPS Glycerol, MOPS Acetate
- PRJEB34578 · SAMEA5972747 · Pseudomonas aerugionsa PAO1 transcriptome cultured in MOPS Glycerol, MOPS Acetate
- PRJEB34578 · SAMEA5972748 · Pseudomonas aerugionsa PAO1 transcriptome cultured in MOPS Glycerol, MOPS Acetate

The title names both carbon sources. Per-record arm assignments came from the source crosswalk.

#### B The shipped statistical interpretation warning

Transcribed from the archived RC.4 build 6 warning capture:

Statistical interpretation. Adjusted P values are exploratory, especially when covariates leave few residual degrees of freedom or study-arm information is missing. Correlation and Fisher-z P values are approximate with tied or sparse data. Normal pooled inference can understate uncertainty with few heterogeneous studies; modified Knapp–Hartung is also assumption-dependent. Shared BioSample checks cannot detect all related studies. A displayed |rho| cutoff is descriptive and does not test that the true effect exceeds that cutoff.
